# γ-Linolenic Acid Induces a Vitamin D Receptor-Independent Mineralization Program by Activating CaMKII–SMAD2/3 Pathway in Calvarial Osteoblasts

**DOI:** 10.64898/2026.06.10.731289

**Authors:** Trisha Biswas, Chen Chongtham, Namaste Kumari, Yatin Saneja, Neeraj Kumar Yadav, Jaswinder Singh Maras, Siddhesh S Kamat, Gopalakrishnan Aneeshkumar Arimbasseri

## Abstract

Vitamin D receptor (VDR) signalling is essential for skeletal mineralisation, and the *vdr /* mouse phenotype has long been attributed principally to disrupted systemic calcium availability. Here we show that VDR is required cell-autonomously to maintain intracellular calcium signalling competence in osteoblasts, and that this requirement can be bypassed metabolically. VDR directly occupies and maintains expression of the osteoblast calcium-handling transcriptome; in its absence, intracellular Ca² dynamics, CaMKII activation, and SMAD2/3 phosphorylation are diminished, arresting osteoblasts before terminal mineralisation. Pharmacological CaMKII inhibition in wild-type cells reproduces this phenotype, establishing CaMKII as an upstream regulator of SMAD2/3, and of mineralisation. Importantly, the omega-6 fatty acid γ-linolenic acid (GLA), identified by serum metabolomics as the circulating mediator of a calcium-independent dietary rescue, restores Ca² flux, CaMKII activation, SMAD2/3 phosphorylation, matrix production and mineralisation in *vdr /* osteoblasts. GLA-mediated rescue is dependent on CaMKII and SMAD2/3 activities. However, this rescue is independent of BMP–SMAD1/5/9 signalling or the early lineage determinants *Runx2* and *Sp7*, demonstrating that terminal mineralisation can be driven via a non-canonical SMAD2/3 route when the canonical BMP arm remains compromised. Consistent with this, SMAD1/5/9 activation is preserved in vivo by paracrine BMP but lost in isolated culture, whereas SMAD2/3 loss persists in both contexts. These findings suggest that the *vdr /* osteoblasts experience a failure of intracellular signalling competence rather than of mineral supply, and identify metabolic modulation of calcium signalling as a route to restoring osteoblast function when vitamin D signalling is impaired.

## Introduction

Bone is a metabolically active organ that regulates mineral homeostasis and systemic physiology (1). Osteoblasts, the bone-forming cells, derive from mesenchymal progenitors and undergo a tightly regulated differentiation programme in which early lineage commitment and late-stage matrix maturation are governed by distinct molecular programmes (2). Disruption of these processes impairs skeletal integrity and contributes to metabolic bone disease (3).

Vitamin D receptor (VDR) signalling is essential for osteoblast maturation, calcium handling and bone integrity, as demonstrated by rickets and impaired mineralisation in *vdr /* mouse models (4). Beyond its role in systemic calcium homeostasis, VDR also regulates intracellular processes, including mitochondrial dynamics, energy metabolism and redox balance (5). Osteoblast differentiation defects in VDR deficiency are stage-specific (6). An *in vitro* study using human osteoblast cell lines showed that calcitriol treatment in the early stages of differentiation, rather than late-stage differentiation enhances mineralization(7).

Two observations challenge the prevailing interpretation of the *vdr /* skeleton that its defects follow from inadequate calcium availability. First, dietary calcium supplementation in mice lacking VDR co-activator binding domain fails to restore skeletal integrity, leaving persistent structural and mineralization deficits (8), implying an osteoblast-intrinsic component that mineral repletion does not reach. Second, the relatively preserved early osteogenic program together with the pronounced defect in late-stage matrix maturation suggests that VDR deficiency affects progression toward the mineralising state in addition to its effects on systemic mineral homeostasis (6). What has been missing is an intracellular mechanism that links VDR activity to the late-stage osteoblast transcriptional programme, and a test of whether that mechanism can be engaged without VDR itself. The importance of resolving this is of general importance, because it determines whether the terminal mineralisation programme is strictly downstream of canonical BMP–SMAD1/5/9–RUNX2 signalling or can it instead be sustained through alternative signalling mechanisms. Here, we identify this signalling mechanism and show that it is metabolically accessible. VDR is required to maintain expression of a subset of calcium-handling genes in osteoblasts; its loss collapses intracellular Ca² dynamics, inactivates CaMKII, and reduces SMAD2/3 phosphorylation, arresting cells short of mineralisation. γ-Linolenic acid (GLA) restores this axis in *vdr /* osteoblasts and rescues matrix production and mineralisation in a CaMKII-dependent manner without restoring BMP–SMAD1/5/9 signalling or the early determinants *Runx2* and *Sp7*. Our findings show that terminal mineralisation can proceed through a non-canonical, SMAD2/3-dependent route and suggest that intracellular signalling competence, rather than receptor occupancy or mineral supply alone, is an important determinant of osteoblast maturation.

## Results

### VDR loss selectively blocks late osteoblast maturation in vivo and in vitro

Given the reduced bone formation and increased skeletal fragility reported in *vdr /* mice (17), we first profiled signalling molecules involved in bone homeostasis in WT and *vdr /* mice using NanoString (Table S1). This analysis aimed to define the molecular alterations associated with this phenotype. Mineralisation-associated transcripts such as Sclerostin (*Sost*), Dentin Matrix Protein 1 (*Dmp1*), Osteocalcin (Bglap), Osteopontin (*Opn*) and Bone Morphogenetic Protein 7 (*Bmp7*) were severely downregulated in *vdr /* bones, whereas osteoglycin (Ogn) was upregulated (Figure 1a; Table S1), consistent with the reduced bone formation and increased fragility reported in these mice.

**Figure 1:**
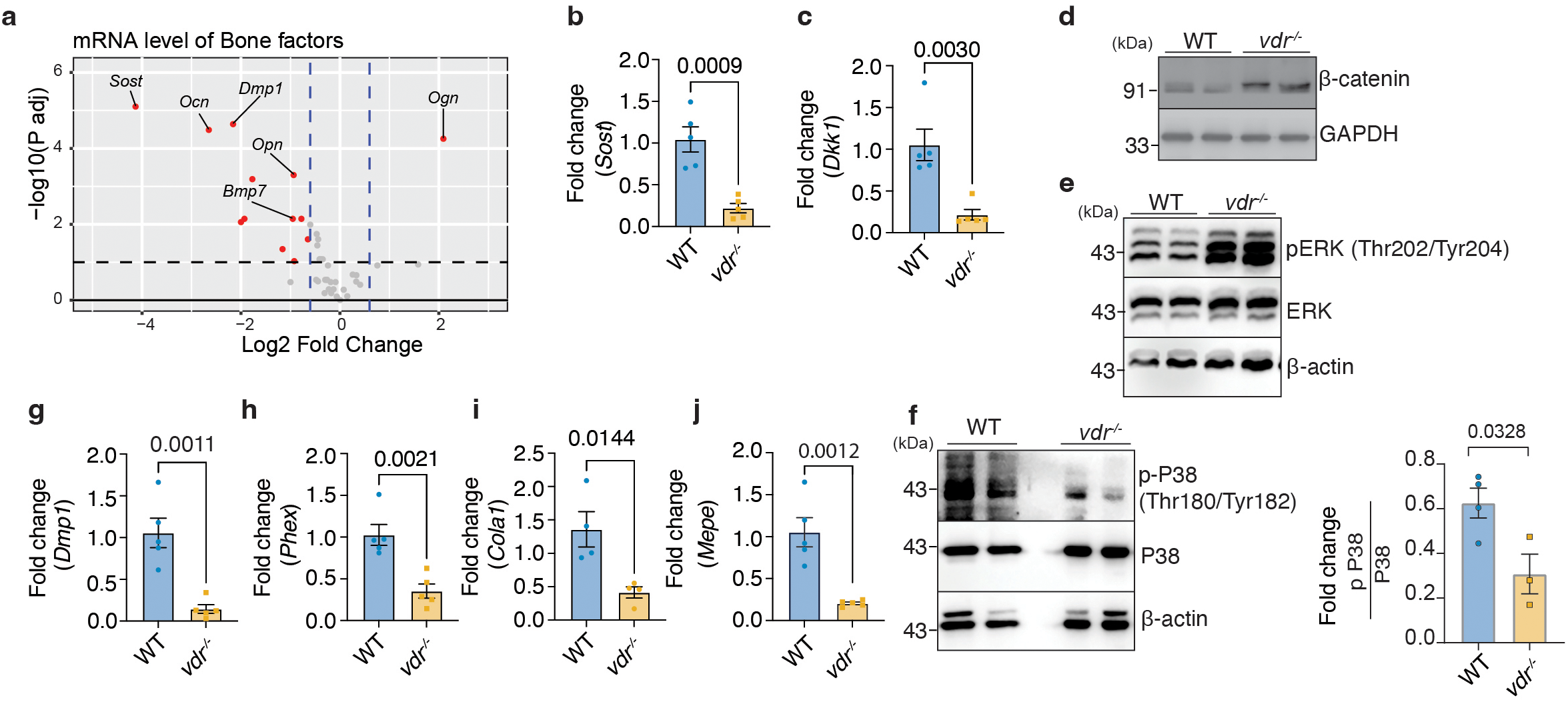
*vdr /* mice exhibit downregulation of late osteogenic signalling through the P38 pathway. (a) Volcano plot showing differential gene expression in *vdr /* osteoblasts relative to WT controls, identified using NanoString technology. Dashed lines indicate thresholds for log2 (fold change) (0.6) and adjusted P value (.1). (b-c) Quantitative PCR analysis of *Sost* (b) and *Dkk1* (c) expression in WT and *vdr /* bones. (d) Immunoblot analysis of β-catenin in WT and *vdr /* bones (n>3). (e-f) Immunoblot analysis of phosphorylated and total ERK1/2 (Thr202/Tyr204) (e) and p38 (Thr180/Tyr182) (f) in WT and *vdr /* bones (n>3) with quantification (right panel) (g-j) Quantitative PCR analysis of osteogenic marker genes *Dmp1* (g), *Phex* (h), *Col1a1* (i) , and *Mepe* (j) in WT and *vdr /* bones. Data are presented as mean ± SEM. Statistical comparisons were performed using unpaired two-tailed Student’s t-test. * denotes P < 0.05, ** P < 0.01, and *** P < 0.001. Individual biological replicates are denoted by dots on the graphs, with the number of biological replicates (*n*) indicated in each graph.

Interestingly, the WNT inhibitors *Sost* and *Dkk1* were reduced (Figures 1b, 1c) (18) and β-catenin was elevated (Figure 1d), indicating heightened WNT signalling. Furthermore, phosphorylation of ERK (Thr202/Tyr204), which supports early differentiation (19), was increased (Figure 1e), suggesting an increase in early differentiation pathways. In contrast, phosphorylated p38 (Thr180/Tyr182), which drives acquisition of a mature mineralising phenotype (20), was significantly reduced (Figure 1f), as were the mature osteocyte and ECM markers *Dmp1, Phex, Mepe* and *Col1a1* (Figures 1g-j). *Runx2* and Osterix (Sp7) were unchanged in vivo (Figure S1a&b), confirming that early specification of osteoblasts is intact. Reduced *Opn* and *Nfatc1*, regulators of osteoclast differentiation and function (21), with unchanged RANKL and OPG protein (Figure S1c-e), indicate that the mineralisation deficit reflects osteoblast maturation failure rather than increased resorption.

To resolve the differentiation stage at which VDR is required, calvarial osteoblasts from WT and *vdr /* mice were differentiated for 20 days and sampled at days 0, 4, 10 and 20, corresponding to early commitment/proliferation, matrix maturation and late mineralization (22). In WT cells, *Dmp1* and *Bglap* rose progressively to a maximum at day 20, while *Phex* and *Col1a1* peaked near day 10. *vdr /* osteoblasts showed markedly reduced expression of all these genes at every time point and failed to mount the characteristic differentiation-associated induction of these genes (Figure 2a), recapitulating the in vivo phenotype. Unlike in vivo, *Runx2* and *Sp7* were reduced in *vdr /* osteoblasts in culture (Figure 2b). Interestingly, alkaline phosphatase staining at day 7 was comparable between genotypes, yet Alizarin Red staining at day 21 was markedly reduced (Figure 2c), indicating that *vdr /* cells retain early osteogenic activity but fail to complete the transition to terminal mineralisation. Consistent with this, phosphorylated p38 was maintained throughout WT differentiation but significantly reduced by day 20 in *vdr /* cultures (Figure 2d), while ERK phosphorylation was unchanged (Figure 2e).

**Figure 2:**
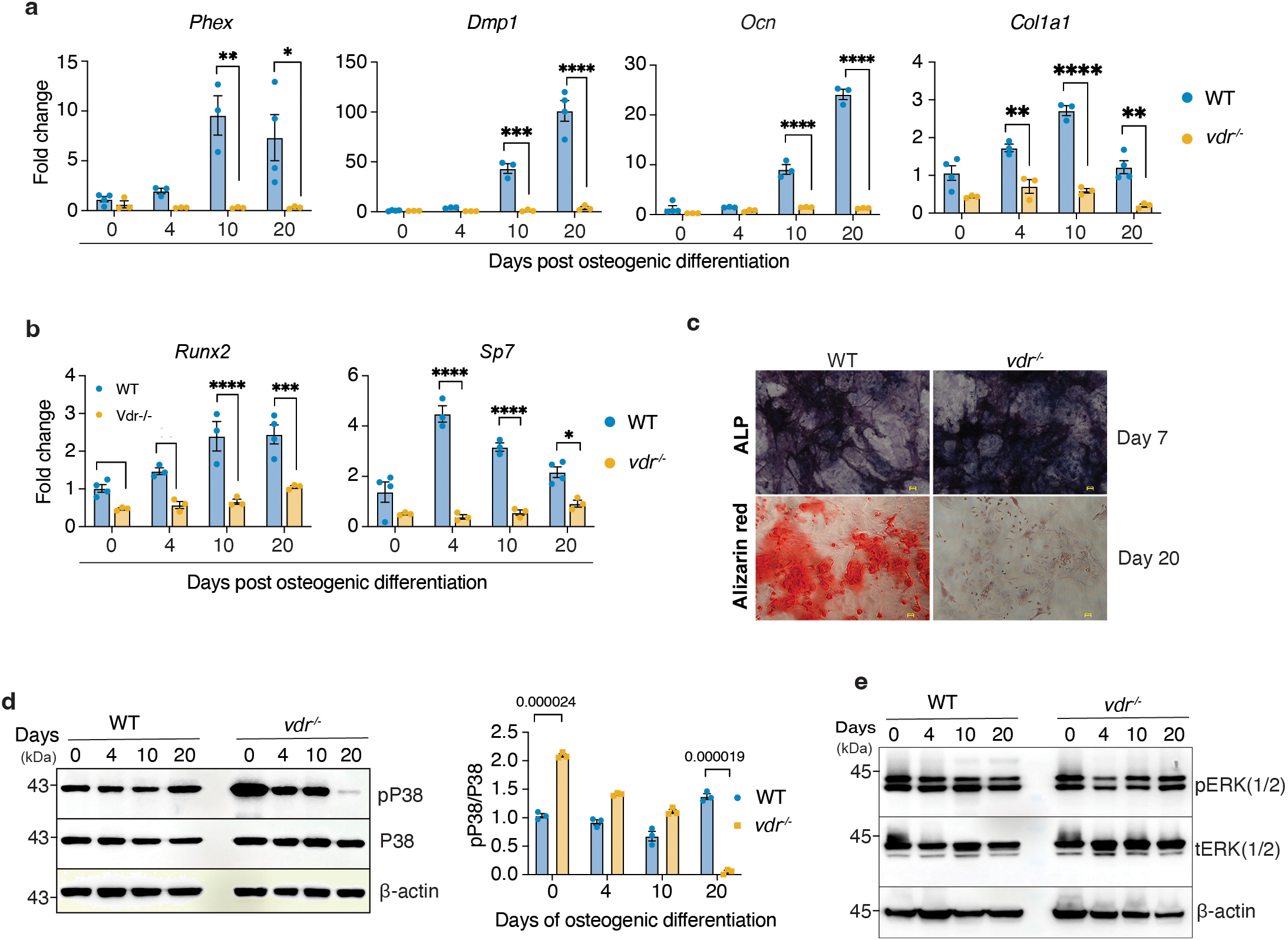
*vdr /*osteoblasts exhibit impaired late differentiation in vitro. (a) Quantitative PCR analysis of osteogenic markers (*Phex, Dmp1,Bglap,* and *Col1a1*) during differentiation (days 0–20) in WT and *vdr /* osteoblasts. (b) Quantitative PCR analysis of *Runx2* and *Osx* (*Sp7*) expression during differentiation in WT and *vdr /* osteoblasts. (c) Representative ALP staining (day 7; upper panel) and Alizarin Red staining (day 20, lower panel) of WT and *vdr /* osteoblasts (scale bar= 24μm). (d-e) Immunoblot analysis of phosphorylated and total p38 with quantification (right) (d) and ERK1/2 (e) during osteogenic differentiation in WT and *vdr /* osteoblasts. Data are presented as mean ± SEM. Statistical comparisons were performed using two- way ANOVA followed by Tukey’s post hoc multiple comparison test. Individual biological replicates are denoted by dots on the graphs, with the number of biological replicates (*n*) indicated in each graph. *P < 0.05, **P < 0.01, ***P < 0.001.

This *in vivo-in vitro* divergence suggests that the defect is not simply due to an accumulation of immature osteoblasts. Despite unchanged early ALP levels, the reduced expression of lineage-determining factors points to a failure to progress toward terminal mineralisation. The reduction in *Runx2* and *Sp7* observed specifically in isolated cultures may reflect the loss of microenvironmental signals, which we examine below. This is also consistent with the coordinated increase in *Vdr* and *Runx2* during osteoblast differentiation (23).

### A cell-autonomous loss of SMAD2/3 activation underlies the maturation block

To understand the defects at the pathway level, we performed transcriptomic profiling across differentiation (days 0, 4, 10, 20). Hierarchical clustering of the top 2000 variable genes yielded four clusters (Figure 3a). Clusters 2 and 3 were enriched in WT cells, with cluster 2 exhibiting early proliferation-associated genes (days 0, 4) and cluster 3 showing late ECM-associated genes (days 10, 20) (Figure S2a). *vdr /* osteoblasts were dominated by cluster 1, which did not respond to differentiation signals and was enriched for chemotaxis and inflammation-related functions, a signature associated with reduced vitamin D function in osteoblasts (6, 24). Time-course trajectory analysis resolved six clusters (Figures 3b, S2b), with late-stage, WT-specific cluster 3 enriched for BMP signalling and ECM genes (Figures 3c, S2b). Transcription factor binding-site analysis across these clusters identified *Smad4* among the principal regulators of the late ECM module, and the inhibitory Smads, *Smad6* and *Smad7*, were upregulated in *vdr /* osteoblasts at this stage (Figure 3d), indicating suppression of BMP and TGF-β output (25) (Figure S3a–c).

**Figure 3.**
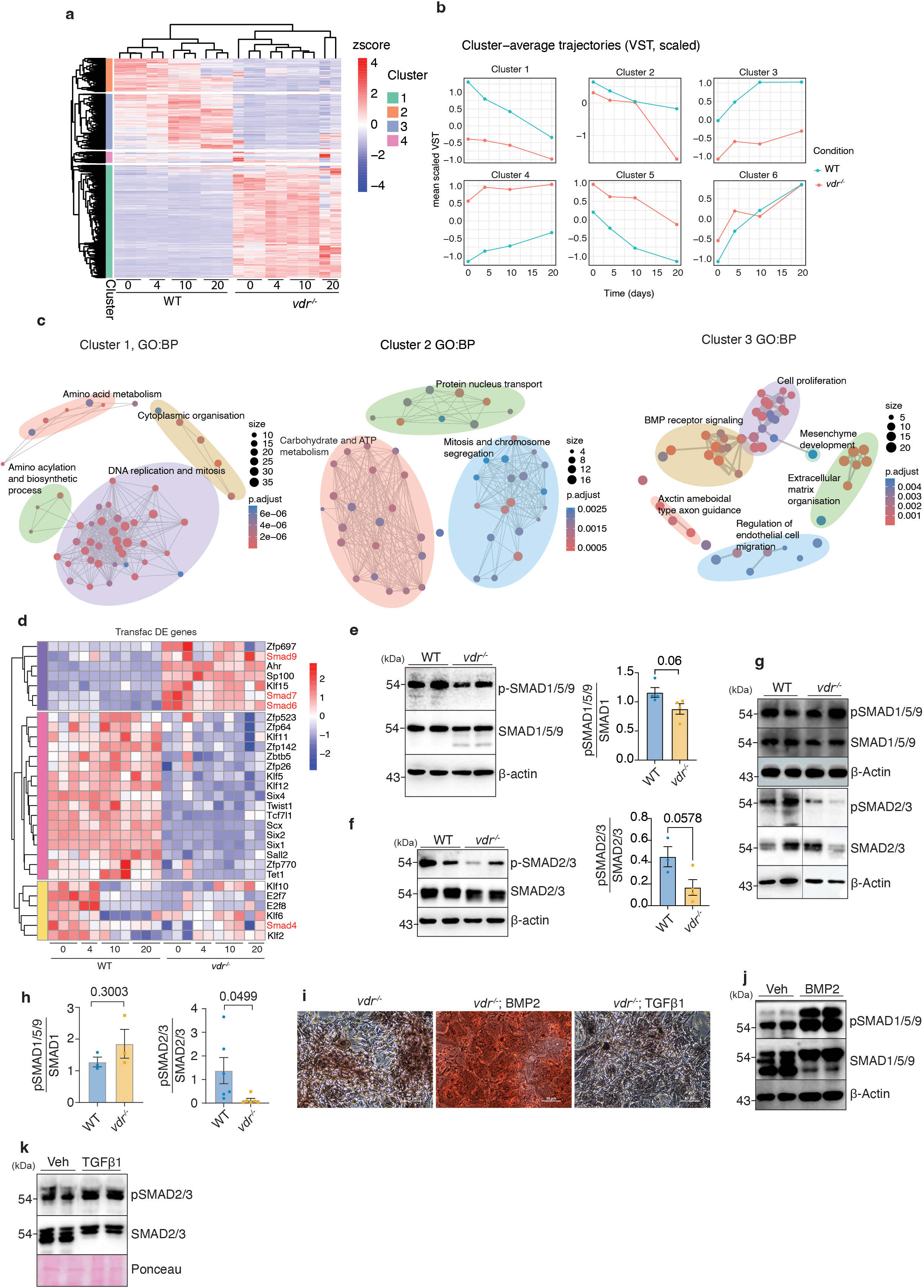
VDR deficiency alters stage-specific SMAD-associated transcriptional programs during osteoblast differentiation. (a) Heatmap showing hierarchical clustering of the top 2000 variable genes across WT and *vdr /* osteoblasts collected at days 0, 4, 10, and 20 of differentiation. Normalised read counts of each gene per condition are mapped. Rows are normalised by z score, and the colour scale shows the z score. 4 different clusters identified are indicated as a colour bar on the left. (b) Time-course trajectory analysis showing six distinct temporal gene expression trajectories during osteoblast differentiation in WT and *vdr /* osteoblasts performed using Deseq2. (c) Gene ontology (GO) enrichment analysis, done using Clusterprofiler, of representative temporal clusters. (d) Heatmap showing expression patterns of transcription factors, whose targets are enriched among the total differentially expressed genes along the course of differentiation, identified using TRANSFAC, across WT and *vdr /* osteoblasts during differentiation. (e) Immunoblot analysis and quantification of phosphorylated and total SMAD1/5/9 in WT and *vdr /* osteoblasts at day 4 of differentiation and, (f) Immunoblot analysis and quantification of phosphorylated and total SMAD2/3 in WT and *vdr /* osteoblasts at day 4 of differentiation (n=4). (g) Immunoblot analysis of phosphorylated and total SMAD1/5/9 and SMAD2/3 in bone lysates from WT and *vdr /* mice bones. Vertical line on the SMAD2/3 panel indicates unrelated lanes spliced out. (h) Densitometric quantification of the blots shown in (g). (i) Alizarin Red staining of *vdr /* osteoblast cultures treated with BMP2 or TGF-β1 during differentiation (scale bar= 50μm). (j) Immunoblot analysis of phosphorylated and total SMAD1/5/9 in *vdr /* osteoblasts following BMP2 treatment (n=3). (k) Immunoblot analysis of phosphorylated and total SMAD2/3 in *vdr /* osteoblasts following TGF-β1 treatment (n=3). Data are presented as mean ± SEM. Statistical analyses were performed using an unpaired two-tailed Student’s t-test. *P < 0.05, **P < 0.01, ***P < 0.001. Individual biological replicates are denoted by dots on the graphs, with the number of biological replicates (*n*) indicated in each graph.

Differential expression at days 0 and 4 supported attenuation of canonical osteogenic SMAD signalling: *Bmp2* was reduced while the extracellular BMP antagonists *Grem1* and *Grem2* were elevated (Figure S3d) (26), and phosphorylation of SMAD1/5/9 (27) was correspondingly reduced at day 4 (Figure 3e). SMAD2/3 phosphorylation (28) was also reduced (Figure 3f) despite increased Tgfb1 and Tgfbr1 transcripts, indicating a decoupling of transcript abundance from signalling output compatible with elevated inhibitory SMAD tone rather than reduced ligand availability.

Interestingly, these two SMAD arms behaved differently *in vivo*. pSMAD1/5/9 was preserved in *vdr /* bones (Figure 3g&h), consistent with paracrine BMP from the bone microenvironment compensating for the autocrine deficit seen in isolated cultures(29); this also accounts for the preservation of *Runx2* and *Sp7 in vivo*, both established SMAD1/5/9 targets. SMAD2/3 phosphorylation, by contrast, was reduced in both contexts (Figures 3g&h), identifying it as a cell-autonomous defect that paracrine signalling cannot rescue. Functional tests confirmed this as exogenous BMP2 restored both SMAD1/5/9 phosphorylation and mineralised nodule formation (Figures 3i&j), showing the BMP arm remains ligand-responsive, whereas TGF-β1 raised SMAD2/3 phosphorylation without efficiently restoring mineralization (Figures 3i&k), consistent with the anti-osteogenic effect of canonical TGF-β signalling on terminal differentiation (30). These results show that VDR is required for autocrine BMP2 activation, which is essential for lineage commitment of ex vivo calvarial cells. On the other hand, though SMAD2/3 is downregulated *in vivo* and *in vitro*, receptor-mediated SMAD2/3 activation in fact prevents mineralization.

### VDR directly maintains the calcium-handling transcriptome and CaMKII activation

To further explore the VDR-dependent and TGFβ-independent SMAD2/3 pathway activation in osteoblasts, we examined publicly available VDR ChIP-seq data from osteoblast cell lines (13). Several calcium channel genes showed strong vitamin D-dependent VDR occupancy at their promoters (Figure S4a), and transcriptomic analysis revealed broad dysregulation of calcium-handling and organellar calcium homeostasis genes in *vdr /* osteoblasts (Figure S4b). Genes showing VDR occupancy were also transcriptionally dysregulated (Figure 4a), suggesting that the calcium-handling machinery is under direct VDR control. Downregulated targets included the mitochondrial uptake components *Mcu* and *Micu1*, the ER release channel *Itpr3* and the plasma membrane channel *Trpv4* (31), while the store-operated components *Stim1*, *Orai1* and *Orai2* (32) were upregulated (Figure 4a).

**Figure 4.**
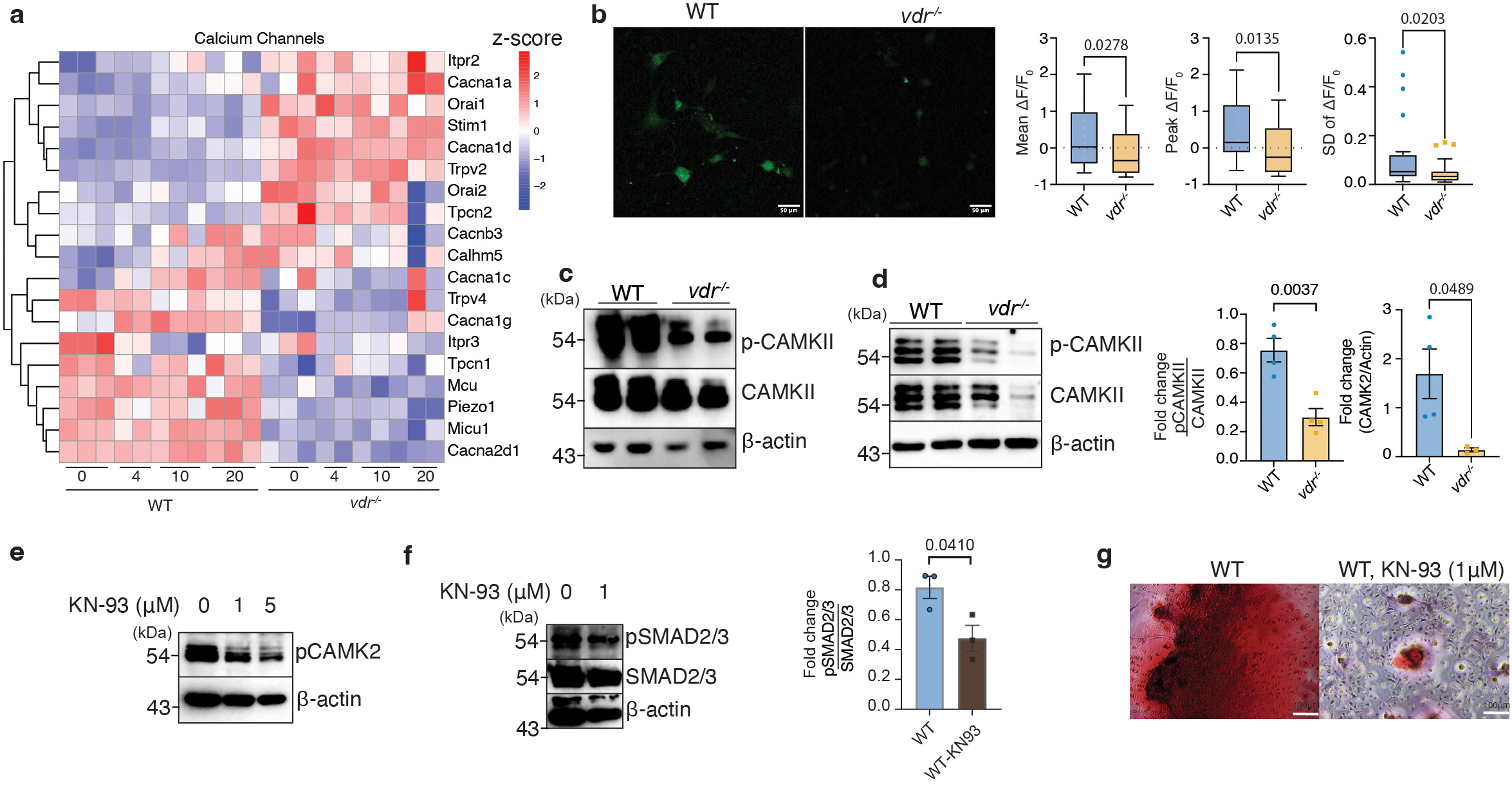
Calcium signalling and CaMKII–SMAD pathway analysis in WT and *vdr /* osteoblasts. (a) Heatmap of calcium channel genes during osteogenic differentiation (days 0, 4, 10, and 20) in WT and *vdr /* osteoblasts. Z scores of Deseq2-normalised read counts are plotted. (b) Live-cell Ca² imaging using Fluo-4 in WT and *vdr /* osteoblasts at day 4 of differentiation. Representative images (scale bar= 50μm) and quantification of mean, peak, and standard deviation of ΔF/F are shown (n=3). (c) Immunoblot analysis of phosphorylated and total CaMKII in bone lysates from WT and *vdr /* mice. (d) Immunoblot analysis of phosphorylated and total CaMKII in WT and *vdr /* osteoblasts at day 4 of differentiation, with quantification (right panel). (e) Immunoblot showing inhibition of CaMKII phosphorylation following treatment with KN-93 (0–5 μM). (f) Immunoblot analysis of phosphorylated and total SMAD2/3 following KN-93 treatment with quantification (right panel). (g) Representative Alizarin Red staining of osteoblast cultures treated with KN-93 (1 μM) (scale bar= 100μm). For figure b, data are presented as box-and-whiskers plots analyzed by a Mann-Whitney test. All other panels are presented as mean ± SEM and analyzed via unpaired two-tailed Student’s t-test where individual biological replicates are denoted by dots on the graphs, with the number of biological replicates (*n*) indicated in each graph. *P < 0.05, **P < 0.01, ***P < 0.001.

In accordance with the transcriptional changes in calcium handling genes, we next asked whether VDR deficiency was associated with an altered basal intracellular Ca² signaling state. Live-cell Fluo-4 imaging showed markedly attenuated basal Ca² oscillations in *vdr /* osteoblasts, with significantly reduced mean, peak and spread of ΔF/F (Figure 4b). Basal Ca² activity is a recognized feature of osteoblasts and can provide a signaling input to Ca² -dependent pathways (33, 34). Thus, these findings indicate that VDR deficiency alters the basal Ca² signaling state of osteoblasts, although they do not establish how the cells respond to a defined external Ca² stimulus. Because intracellular Ca² handling is closely integrated with mitochondrial function, we next asked whether the altered Ca² state was accompanied by broader mitochondrial dysfunction. Seahorse analysis revealed reduced basal respiration and spare respiratory capacity without major changes in glycolysis (Figure S5a–d), and mitochondrial superoxide was elevated (Figure S5e,f), indicating that mitochondrial function was also affected in *vdr /* osteoblasts. However, the mitochondria-targeted antioxidant MitoTEMPO only partially restored nodule formation (Figure S5g) and failed to upregulate mineralisation markers other than *Dmp1* (Figure S5h), suggesting that the increased mitochondrial ROS is not the primary defect in the *vdr /* osteoblasts.

Because intracellular Ca² dynamics are the major upstream input to CaMKII activation (34), we next assessed CaMKII activation. Total CaMKII was largely preserved, but its phosphorylation was markedly reduced in *vdr /* bones (Figure 4c), and differentiated *vdr /* osteoblasts showed reduced CaMKII protein and phosphorylation *in vitro* (Figure 4d). VDR loss therefore leads to direct loss of calcium-handling gene expression, collapse of Ca² dynamics, and failure of CaMKII activation.

### CaMKII is required for SMAD2/3 activation and mineralisation

Since we observed defective SMAD2/3 signaling and CaMKII activation, we tested whether these two signaling hubs are connected. We inhibited it pharmacologically using KN-93 in wild-type osteoblasts (Figure 4e). KN-93 reduced SMAD2/3 phosphorylation without affecting total SMAD2/3 (Figure 4f), establishing CaMKII as an upstream regulator of SMAD signalling in osteoblasts. Functionally, CaMKII inhibition markedly impaired mineralised nodule formation in WT cultures (Figure 4g). CaMKII activity is thus required both for SMAD2/3 signalling and for mineralisation, and its loss is sufficient to reproduce the *vdr /* phenotype in cells with intact VDR. Taken together, these results identify a TGF-β-independent route to SMAD2/3 activation and its role in osteoblast mineralisation.

### A milk-based diet rescues the skeleton independent of calcium and identifies GLA as a candidate mediator of the dietary response

*vdr /* mice do not develop rickets during suckling, when milk is the primary nutritional source(17, 35), and we previously showed that milk-based diets (MBD) rescue metabolic defects in these mice without restoring serum calcium (36, 37). Weaning *vdr /* mice onto MBD at 3 weeks (Figure 5a) restored body size, weight and tibial length by 7 weeks (Figures 5b, S6a) and normalised the widened growth plates and expanded hypertrophic chondrocyte layers characteristic of rickets (Figure 5c). PTH remained elevated, and circulating vitamin D remained altered (Figures S6b, S6c), suggesting that the skeletal rescue occurs without restoring VDR-dependent calcium homeostasis. MBD increased *Dmp1, Phex, Mepe, Sost* and *Col1a1* (Figure 5d) and restored phospho-p38 (Figure 5e)., Importantly, MBD restored CaMKII phosphorylation (Figure 5f) and pSMAD2/3 (Figure 5g) in *vdr /* bones, while SMAD1/5/9 phosphorylation remained comparatively unchanged (Figure S6d), indicating a selective *in vivo* recovery of the CaMKII-SMAD2/3 axis identified above.

**Figure 5.**
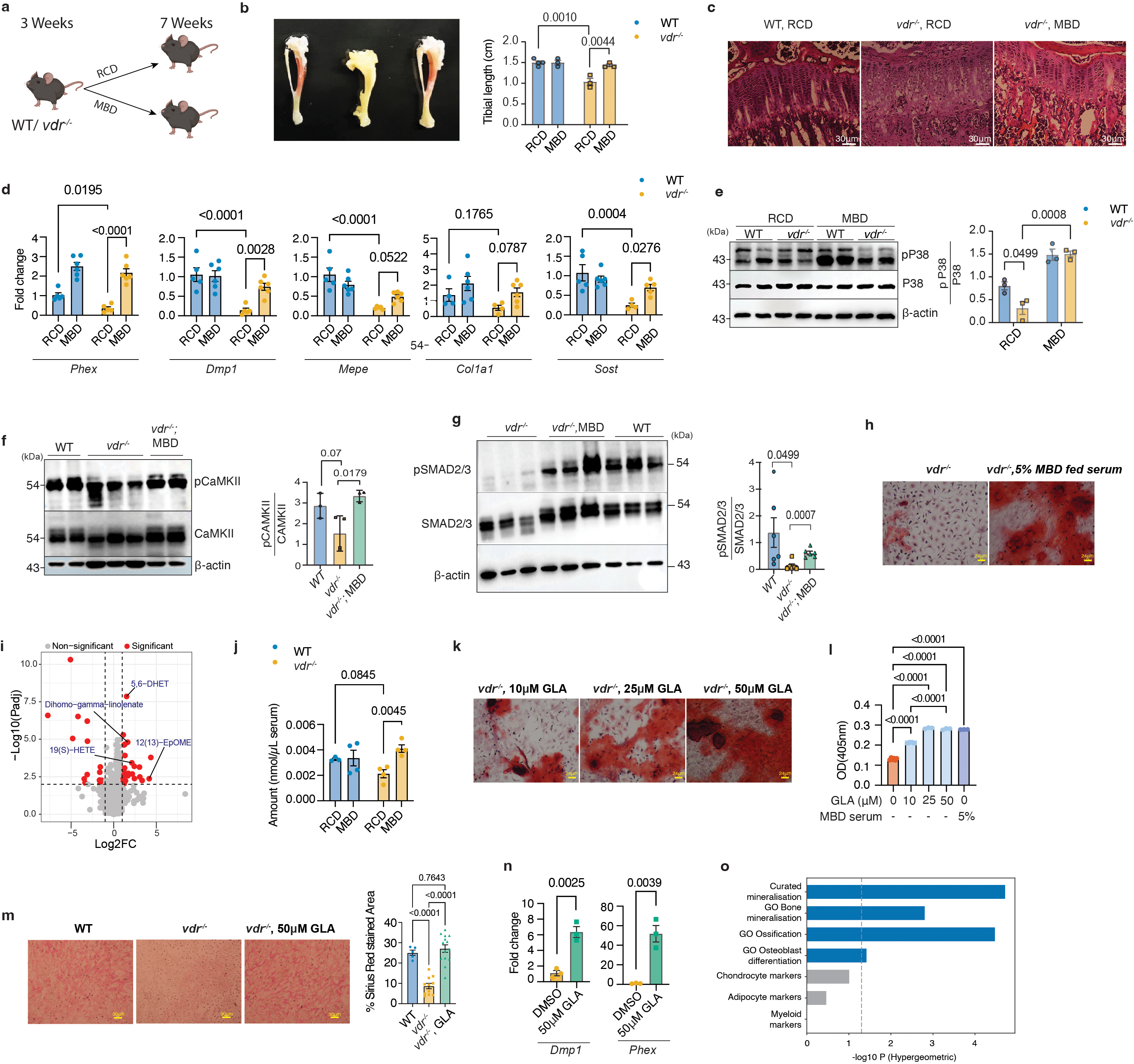
A milk-based diet restores skeletal abnormalities and promotes osteoblast maturation in *vdr /*mice. (a) Schematic representation of the experimental design showing dietary intervention of *vdr /* mice with chow diet or milk-based diet (MBD) following weaning. (b) Representative images of mice and tibiae from WT, chow-fed *vdr /* (left panel), and MBD-fed *vdr /* groups, along with quantification of tibial length (right panel). (c) Representative H&E-stained sections of tibial growth plates from WT, chow-fed *vdr /* , and MBD-fed *vdr /* mice. Magnification: 20X. Scale bars indicate 30μm. (d) Quantitative PCR analysis of osteogenic and osteocyte-associated genes (*Phex, Dmp1, Mepe, Col1a1* and *Sost*) in bone tissue from the indicated groups (n>5). (e) Immunoblot analysis of phosphorylated and total p38 MAPK in bone lysates from chow-fed and milk-based diet (MBD)-fed WT and *vdr /* mice. Quantification is given on the right panel. (f) Immunoblot analysis of phosphorylated and total CaMKII in bone lysates from WT, chow-fed *vdr /* , and MBD-fed *vdr /* mice. Quantification is given on the right panel. (g) Immunoblot analysis of phosphorylated and total SMAD2/3 in bone lysates from WT, chow-fed *vdr /* , and MBD-fed *vdr /* mice. Quantification is given on the right panel. (h) Representative Alizarin Red staining of *vdr /* osteoblast cultures treated with serum from chow-fed or MBD-fed mice (scale bar= 24μm). (i) Volcano plot showing differentially abundant serum metabolites between chow-fed and MBD-fed conditions. Dashed lines indicate the significance thresholds used to define differential metabolites (FDR < 0.1 and |log FC| > 0.5). (j) Quantification of γ-linolenic acid (GLA) levels in serum from the indicated groups. (k) Representative Alizarin Red staining of *vdr /* osteoblast cultures treated with increasing concentrations of GLA (scale bar= 24μm). (l) Quantification of mineralisation by Alizarin Red staining in *vdr /* osteoblasts treated with increasing concentrations of GLA. (m) Representative Sirius Red staining of WT, *vdr /* , and GLA-treated *vdr /* osteoblast cultures (scale bar= 30μm) with quantification (right panel). (n) Quantitative PCR analysis of *Dmp1* and *Phex* expression in *vdr /* osteoblasts following GLA treatment. (o) Gene-set enrichment analysis of the GLA rescue gene set, defined as genes significantly downregulated in *vdr* / osteoblasts relative to WT and significantly upregulated following GLA treatment. Enrichment is shown for curated mineralisation and osteoblast-associated gene sets and for unrelated lineage-associated gene sets, including chondrocyte, adipocyte, and myeloid markers. The x-axis represents −log10 *P* values; the dashed line indicates the nominal *P* = 0.05 significance threshold. Data are presented as mean ± SEM; For multi-group comparisons (panels b, d, e, j), data were analyzed by two-way ANOVA followed by Tukey’s post-hoc test. For panels f, g, h, l, m, n, comparisons were analyzed using an unpaired two-tailed Student’s t-test. *P < 0.05, **P < 0.01, ***P < 0.001. Individual biological replicates are denoted by dots on the graphs, with the number of biological replicates (*n*) indicated in each graph.

Serum from MBD-fed mice enhanced mineralised nodule formation in *vdr /* osteoblast cultures relative to chow-fed serum (Figure 5h), indicating a circulating metabolite could be responsible for restoration of mineralisation. Untargeted metabolomics identified elevated derivatives of the omega-6 fatty acid γ-linolenic acid (GLA) in MBD serum (Figure 5i), and targeted metabolomics confirmed elevated GLA in MBD-fed *vdr /* serum (Figure 5j).

### GLA restores mineralisation through CaMKII-dependent SMAD2/3 activation

Exogenous GLA restored mineralised nodule formation in *vdr /* osteoblast cultures dose-dependently, with maximal efficacy at 50 μM (Figures 5k&l). Interestingly, at this concentration, GLA exerted opposing effects depending on VDR status, reducing mineralisation in WT osteoblasts while restoring mineralisation in *vdr* / cultures (Figure S6e). A downstream metabolite of GLA, arachidonic acid, failed to rescue mineralisation even at 50 μM (Figure S6f), establishing the specificity of GLA in restoring mineralisation. GLA also restored collagen matrix accumulation, which was markedly reduced in *vdr /* cultures (Figure 5m), indicating recovery of matrix production preceding mineralisation, while reducing collagen matrix accumulation in WT cells (Figure S6e). GLA also selectively induced the late markers *Dmp1* (5.2-fold) and *Phex* (50-fold) (Figure 5n) together with elevated phospho-p38 at day 20 in *vdr /* osteoblasts (Figure S6g&h). Transcriptome analysis showed that the GLA rescue group (the genes that were significantly downregulated in *vdr /* osteoblasts compared with WT, but upregulated when GLA was added), was enriched for gene sets associated with mineralisation, while the unrelated gene sets, such as chondrocyte, adipocyte and myeloid markers, were unaffected (Figure 5o & S6i).

We next asked whether GLA engages the Ca² –CaMKII signalling defect identified in *vdr* / osteoblasts. Importantly, GLA markedly increased both total CaMKII and CaMKII phosphorylation in *vdr /* osteoblasts (Figure 6a&S6j). Live-cell Fluo-4 imaging revealed restored basal intracellular Ca² dynamics, with increased mean and peak ΔF/F and greater temporal variability relative to untreated *vdr /* cells over a 72–second window (Figures 6b & c). GLA also reduced mitochondrial ROS (Figure 6d), consistent with restored mitochondrial redox homeostasis. GLA increased SMAD2/3 phosphorylation in *vdr /* osteoblasts, whereas SMAD1/5/9 phosphorylation remained unchanged (Figures 6e,f,& S6j). Accordingly, *Runx2* expression was not significantly altered by GLA (Figure 6g).

**Figure 6:**
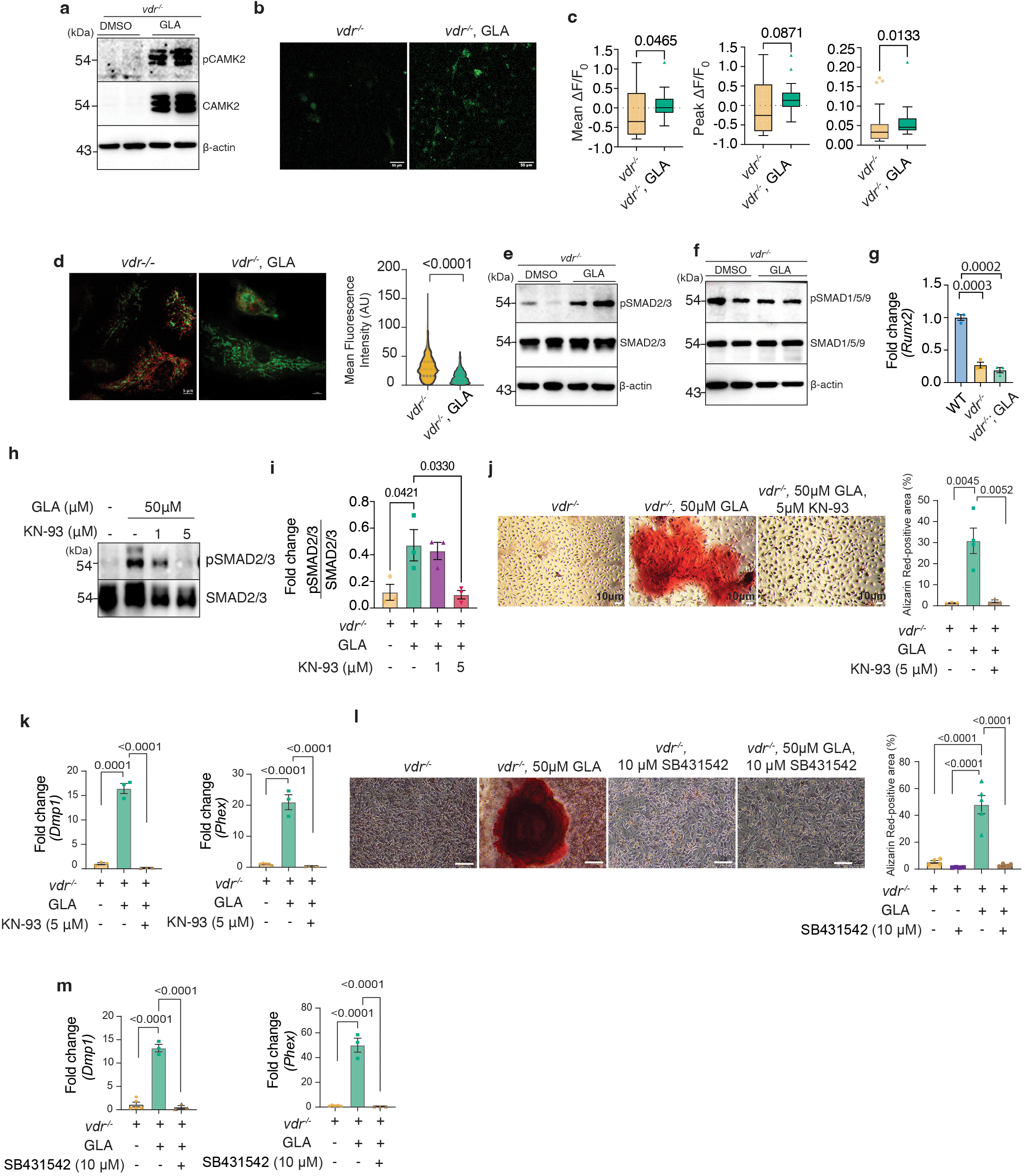
GLA restores Ca² signalling and activates CaMKII–SMAD2/3 signalling in *vdr /* osteoblasts. (a) Immunoblot analysis of phosphorylated and total CaMKII in *vdr /* osteoblasts treated with DMSO or GLA. (b) Representative Fluo-4 Ca² imaging of *vdr /* osteoblasts treated with or without GLA (scale bar= 50 μm). (c) Quantification of mean, peak, and standard deviation of ΔF/F from Ca² imaging traces in *vdr /* osteoblasts under the indicated conditions (n=3). (d) Representative images of mitochondrial ROS in *vdr /* osteoblasts ± GLA, day 7 of osteogenic differentiation (scale bar=5μm). Median MitoSOX fluorescence intensity within mitochondrial ROIs from 3 independent biological replicates. (e-f) Immunoblot analysis of phosphorylated and total SMAD2/3 (e) and SMAD1/5/9 (f) in *vdr /* osteoblasts treated with DMSO or GLA. (g) Quantitative PCR analysis of *Runx2* expression in *vdr /* osteoblasts following GLA treatment. (h) Immunoblot and (i) quantification of phosphorylated SMAD2/3 in *vdr /* osteoblasts treated with GLA and KN-93 at indicated concentrations. (j) Representative Alizarin Red staining of *vdr /* osteoblasts treated with GLA (50 μM) with or without KN-93 (5 μM) (scale bar=10μm) and quantification of Alizarin Red- positive area (%), measured by ImageJ analysis of the stained area. (k) Expression of *Dmp1* and *Phex* in *vdr*−/− osteoblasts treated with GLA alone or GLA in combination with KN-93. (l) Mineralisation in *vdr*−/− osteoblasts treated with GLA alone or GLA in combination with the SMAD2/3 phosphorylation inhibitor SB431542 (scale bar=100μm) and quantification of Alizarin Red-positive area (%), measured by ImageJ analysis of the stained area. (m) Expression of *Dmp1* and *Phex* following treatment with GLA alone or GLA in combination with SB431542. For figure c and d, data are shown as box-and-whiskers plots and violin plots (median ± quartiles) respectively and analyzed by a Mann-Whitney test. All other figures are presented as mean ± SEM and analyzed via one-way anova where individual biological replicates are denoted by dots on the graphs, with the number of biological replicates (*n*) indicated in each graph. *P < 0.05, **P < 0.01, ***P < 0.001.

Notably, GLA also increased SMAD2/3 phosphorylation in WT osteoblasts (Figure S6k), despite reducing mineralisation in these cells. Thus, increased SMAD2/3 phosphorylation alone was not sufficient to determine the mineralisation response. Rather, these findings indicate that the mineralisation response to GLA depends on the signalling context and level of SMAD2/3 activity established by VDR status. The results so far indicate that GLA restores mineralisation specifically in *vdr* / through a Ca² –CaMKII– SMAD2/3-dependent mechanism rather than through reactivation of the canonical BMP– SMAD1/5/9–RUNX2 programme. Finally, we tested whether CaMKII is required for this effect. KN-93 markedly reduced CaMKII phosphorylation (Figure S6l) and suppressed the GLA-induced increase in SMAD2/3 phosphorylation (Figures 6h&i), and co-treatment with GLA and KN-93 markedly reduced mineralisation and expression of *Dmp1* and *Phex* (Figure 6j&k). These results strongly suggest that CaMKII is a critical mediator of GLA-induced SMAD2/3 activation and mineralisation rescue in *vdr* / osteoblasts. Co-treatment of the *vdr /* osteoblasts with SB431542, an inhibitor of SMAD2/3 phosphorylation and GLA, abolished the SMAD2/3 phosphorylation (Figure S6m&n) and mineralisation induced by GLA (Figure 6l). qRT-PCR analyses showed reduced expression of *Dmp1* and *Phex* when GLA treated *vdr /* osteoblasts were subjected to SB431542 co-treatment (Figure 6m). These findings support a requirement for SMAD2/3 signaling in the GLA-mediated restoration of mineralization.

## Discussion

While the *vdr /* mouse model has existed for the past 30 years, the finding reported here highlights an unexpected functional connection between the vitamin D receptor, CaMKII and SMAD signalling pathways in osteoblasts. Our results argue that the limiting determinant of *vdr /* osteoblast maturation is intracellular signalling competence rather than simply a consequence of receptor occupancy or mineral supply, and reveal the functional flexibility of the SMAD network. Two findings reported here establish this flexible signalling framework in osteoblasts. First, terminal osteoblast mineralisation can be restored by GLA despite persistent suppression of canonical BMP– SMAD1/5/9–RUNX2 signalling. GLA restores the full late-stage mineralisation module while SMAD1/5/9 phosphorylation remains suppressed, and *Runx2* and *Sp7* remain at lower levels. Transcriptome analysis confirms that GLA activates the mineralisation gene module without restoring SMAD1/5/9 targets. Second, a dietary lipid can substitute for the intracellular signalling output of a nuclear receptor, restoring the Ca² –CaMKII– SMAD2/3 axis in cells that lack VDR entirely, indicating that SMAD2/3 can sustain the late-stage mineralisation module even when the canonical BMP2-*Runx2* arm is impaired.

The behaviour of the two SMAD arms supports this interpretation. SMAD1/5/9, the BMP target and an essential determinant of osteogenesis (27), is downregulated in vitro but unaffected in vivo, consistent with paracrine BMP from osteoclasts, marrow stromal cells and endothelial cells sustaining signalling in intact bone while isolated cultures depend on autocrine ligand (29). This also explains the *in vivo-in vitro* divergence in *Runx2* and *Sp7*, both established SMAD1/5/9 targets (38). SMAD2/3, by contrast, is reduced in both contexts, marking it as the cell-autonomous lesion. The functional tests reveal a dichotomy between the two arms. BMP2 supplementation restored mineralisation, confirming the BMP arm remains competent when ligand is supplied, whereas TGF-β1 raised SMAD2/3 phosphorylation without restoring mineralisation, consistent with the stage-dependent, anti-osteogenic action of canonical TGF-β on terminal differentiation (30). Thus, TGF-β1–driven SMAD2/3 phosphorylation is not sufficient to restore late mineralisation, while inhibition of SMAD2/3 prevents mineralisation, suggesting that the functional consequence of SMAD2/3 activation depends on signalling context. We conclude that receptor-driven phosphorylation of SMAD2/3 does not translate to functionally productive signalling.

Interestingly, a CaMKII-dependent phosphorylation of SMAD2/3 is accountable for that discrepancy. Previous work established that CaMKII is required for osteoblast differentiation and bone formation, with inhibition reducing osteogenic gene expression and mineralization (34, 39), that it modulates SMAD phosphorylation and signalling output across different systems (40, 41), and that disrupting Ca² –CaMKII signalling impairs SMAD activation in genetic bone disease (41). Here, *vdr /* osteoblasts show impaired Ca² dynamics and diminished CaMKII activation with reduced SMAD2/3 phosphorylation. Pharmacological CaMKII inhibition in wild-type cells reproduces both the signalling and the mineralisation phenotype, placing CaMKII upstream of SMAD2/3 in this response and identifying a CaMKII–dependent route contributing to SMAD2/3 activation. Consistent with this, inhibition of SMAD2/3 phosphorylation with SB431542 abolished the mineralisation rescue produced by GLA. Together with the KN-93 experiments, these findings establish a functional requirement for both CaMKII activity and SMAD2/3 signalling in the GLA response. The precise relationship between CaMKII and ALK4/5/7-mediated SMAD2/3 signalling remains unresolved; our data do not establish whether CaMKII acts upstream of receptor-mediated SMAD2/3 activation, in parallel with it, or modulates its downstream output. The extent to which SMAD2/3 activity is strictly required for the differentiation phenotype will need direct genetic interrogation. We suggest that the upstream defect is transcriptional; ChIP-seq and RNA-seq together show that a major subset of calcium transport genes are under VDR control in osteoblasts, and that VDR loss produces a coordinated collapse of the canonical Ca² signalling architecture. Principal entry and release channels, the L-type and T-type voltage-gated channels *Cacna1c* and *Cacna1g*, the mechanosensitive channels *Piezo1* and *Trpv4*, and the ER release channel IP R3 (*Itpr3*), are downregulated, likely impairing the cytosolic transients required for CaMKII Thr286 autophosphorylation. Dysregulation of the mitochondrial uniporter complex (*Mcu, Micu1*) would further distort transient kinetics. Upregulation of the SOCE components STIM1, ORAI1 and ORAI2, the ER isoform IP R2 (Itpr2), the atypical voltage-gated channels Cav2.1 (*Cacna1*a) and Cav1.3 (*Cacna1d*), and the high-threshold channel TRPV2 appears maladaptive, as it does not restore CaMKII activation.

Interestingly, at 50 μM, the concentration that maximally restored mineralisation in *vdr* / osteoblasts GLA exerted an opposing effect in VDR-replete cells. Notably, GLA increased SMAD2/3 phosphorylation in both WT and *vdr* / osteoblasts despite these opposite effects on mineralisation, indicating that increased SMAD2/3 phosphorylation alone is not sufficient to determine the mineralisation response. Rather, the effect of GLA on mineralisation appears to depend on VDR status and the associated basal SMAD2/3 signalling state: GLA increased SMAD2/3 signalling in WT cells, where basal signalling was preserved, while restoring diminished SMAD2/3 signalling in *vdr* / cells. How GLA re-engages this axis remains open. The failure of arachidonic acid, a downstream metabolite of GLA, to restore mineralisation indicates that GLA itself, rather than its metabolites, mediates the effect. This also rules out a general effect of PUFA on membrane fluidity. Interestingly, GLA is a ligand for nuclear receptors including PPAR- γ, PPAR-δ and the VDR heterodimeric partner RXR (42–44), and may therefore restore calcium-handling gene expression through alternative nuclear receptors; this remains to be tested directly. The present findings also raise the possibility that GLA does not simply restore SMAD2/3 phosphorylation, but engages SMAD2/3 in a signalling context that is permissive for late mineralisation. The fact that TGF-β1 increased SMAD2/3 phosphorylation without rescuing mineralisation, whereas GLA-associated SMAD2/3 activation was accompanied by restoration of Ca² –CaMKII signalling and mineralisation, supports this distinction.

Testing whether GLA acts through this route, how CaMKII intersects with ALK4/5/7– SMAD2/3 signalling, and whether the CaMKII–SMAD2/3 axis is engageable in mineralisation defects beyond VDR deficiency, are the immediate priorities to take forward.

## Conclusion

Our study defines a framework in which VDR sustains osteoblast maturation by maintaining intracellular Ca² dynamics and CaMKII activity, thereby enabling SMAD2/3-dependent matrix production. In the absence of VDR, disruption of this axis, along with the BMP2-SMAD1/5/9-RUNX2 axis, arrests maturation despite available TGF-β ligand and intact early differentiation programmes. Metabolic factors such as GLA bypass this defect by restoring Ca² –CaMKII signalling and reactivating SMAD2/3–dependent transcription, without restoring canonical BMP–SMAD1/5/9– RUNX2 signalling. Intracellular signalling competence therefore emerges as a critical and metabolically tractable determinant of osteoblast differentiation, and metabolic modulation of calcium–dependent signalling offers a route to restoring osteoblast function where vitamin D signalling is impaired.

## Experimental Procedures

### Animals and dietary intervention

*vdr /* mice (Stock No. 006133, B6.129S4-Vdrtm1Mbd/J; Jackson Laboratory, Bar Harbor, ME, USA) were maintained under SPF conditions in individually ventilated cages at the National Institute of Immunology, New Delhi. Genotyping was performed on each litter, and *vdr /* mice and WT littermate controls were selected to minimise genetic variability. Researchers were not blinded, and mice were group-housed. At 3 weeks of age, mice were weaned onto standard rodent chow (Altromin 1324) or milk-based diet (MBD; Lactogen 2, Nestlé) ad libitum for 4 weeks. Both sexes were included as no sex-specific differences were observed. Mice were euthanised at 7 weeks and tissues collected for downstream analyses. All procedures were approved by the Institutional Animal Ethics Committee of the National Institute of Immunology and conducted in accordance with institutional guidelines.

### Tibial bone isolation and processing

Hind limb bones (one femur and one tibia per animal) were isolated from 7-week-old wild-type and *vdr /* mice. After removal of muscles and other surrounding soft tissues, the bone surfaces were gently wiped with sterile gauze to remove residual adherent tissue, bones were washed twice in ice-cold phosphate-buffered saline (PBS). Bone marrow was removed by centrifugation (10,000 × g, 1 min) using an 18-gauge needle-punctured 0.5 mL tube nested within a 1.5 mL tube. Marrow pellets were discarded, and flushed bones were snap-frozen in liquid nitrogen and stored at −80°C for subsequent RNA and protein extraction.

### Histology and cryosectioning

Tibiae were fixed in 4% PFA (overnight, 4°C), decalcified in 20% EDTA (pH 7.4, 2 weeks), embedded in OCT, and cryosectioned at 10 μm (Thermo Scientific HM525 NX). Sections were stained with H&E (Sigma-Aldrich) and imaged at 20× (Olympus inverted microscope). Four images per sample from non-overlapping fields were acquired (n > 3).

### Primary osteoblast culture

Primary calvarial osteoblasts were isolated from 2-week-old pups as described (9). Calvariae were dissected, stripped of connective tissue, minced, and sequentially digested with 0.25% trypsin (Gibco; 10 min) and collagenase II (Gibco; 60 min, 37°C with agitation). Bone chips (20–30 per T25 flask) were plated in α-MEM (Gibco) with 10% FBS (Gibco) and 1% penicillin–streptomycin (Gibco); medium was replaced every 2–3 days. Passages 2–4 were used.

### Osteogenic differentiation and experimental treatments

Osteogenic differentiation was induced at 90–100% confluence in α-MEM supplemented with 10% FBS, 1% penicillin–streptomycin, 10 mM β-glycerophosphate (HiMedia), and 50 μg/mL ascorbic acid (HiMedia), refreshed every 2–3 days for up to 20 days. For stimulation experiments, recombinant BMP2 (100 ng/mL; Thermo Fisher) or TGF-β1 (1 ng/mL; BioLegend) was added to differentiation cultures; cells were harvested at day 4 for signalling analyses or maintained until day 20 for mineralisation assays. GLA (Sigma) or arachidonic acid (Sigma) was prepared in DMSO, aliquoted to minimize repeated freeze–thaw cycles, stored in -20 degrees, and protected from light. Working solutions were freshly prepared immediately before addition to cultures and added at 10, 25, or 50 μM; controls received equivalent DMSO. CaMKII inhibitor KN-93 (Sigma) was prepared as a 5 mg/mL (8.35 mM) stock in Milli-Q water and used at 1 or 5 μM. SB431542 (Sigma) was prepared as a 10 mM stock in DMSO and used at 10 μM. Unless otherwise specified, inhibitor treatments were maintained throughout differentiation until the indicated endpoint. MitoTEMPO (Sigma; 5 μM) was applied over defined temporal windows (days 0–10, 10–20, or 0–20). Serum from chow- or MBD-fed mice was added at 5% v/v. Cells were harvested at day 4 for signalling analyses or day 20 for mineralisation assays unless otherwise specified.

### RNA isolation and quantitative real-time PCR

Total RNA was isolated from tibial bone tissue and primary osteoblasts. Bone samples were pulverised in liquid nitrogen, homogenised in RNAiso Plus (Takara; 1 mL/100 mg tissue), and processed by chloroform extraction and isopropanol precipitation. RNA was washed with 70% ethanol and resuspended in nuclease-free water. RNA concentration and purity were assessed using a Multiskan Sky spectrophotometer (Thermo Fisher); samples with A260/A280 ≈ 2 were used for analysis. RNA from primary osteoblasts was isolated using the NucleoSpin kit (Macherey-Nagel) according to the manufacturer’s instructions. cDNA was synthesised using PrimeScript (Takara), and qPCR was performed with SYBR Premix Ex Taq (Takara) on a QuantStudio 6 Flex system (Thermo Fisher). Gene expression was normalised to β-actin, and the fold changes were calculated using the 2−ΔΔCt method. All reactions were performed in technical triplicate (primers listed in Table S2).

### NanoString gene expression analysis

A custom NanoString panel targeting 45 signaling molecules (including 20 osteokines) and housekeeping genes (HPRT1, HMBS, Polr2b; Table S1) was hybridised with 100 ng RNA at 65°C overnight and processed on the nCounter Analyzer. Raw counts were normalised in nSolver v4.0 (background correction, positive control normalisation, housekeeping gene normalisation) and log -transformed (10).

### RNA sequencing and analysis

RNA-seq libraries were prepared from WT and *vdr /* calvarial osteoblasts at days 0, 4, 10, and 20 of differentiation, and from GLA-treated *vdr /* osteoblasts at days 4, 10, and 20 (n = 3 per group; RQN ≥ 7). rRNA depletion and strand-specific library preparation were performed using QIAseq FastSelect kits (Qiagen) and sequenced on an Illumina NextSeq 2000 (50 bp paired-end). Reads were trimmed (Trimgalore v0.6.10), aligned to GRCm38/mm10 (STAR v2.7.11b), and quantified with featureCounts (Subread v2.0.8); Ensembl IDs were mapped to MGI symbols (biomaRt v2.56). Differential expression was performed in R (v4.3.0) with DESeq2 (v1.40)(11) using a single-factor design encoding genotype–timepoint combinations; genes with mean counts < 100 were excluded. An outlier sample (KO_20D replicate 3, identified by PCA) was excluded from the GLA experiment. DEGs: adjusted p < 0.05 (Benjamini–Hochberg) and |log FC| > 1. Genotype-by-time interactions were identified by LRT using natural cubic splines (df = 3); significant genes were hierarchically clustered (Euclidean, complete linkage) into six trajectory clusters. Unsupervised clustering used the top 2,000 high-variance genes (k = 4; pheatmap v1.0.12). GO enrichment used clusterProfiler (v4.8)(12) against all expressed genes (adjusted p ≤ 0.05; enrichplot v1.20). TF binding motifs were analysed using TRANSFAC (BIOBASE) on promoters (−5 kb to +1 kb).

### Public ChIP-seq analysis

Publicly available VDR and RXR ChIP-seq data were obtained from GEO (GSE51515) (13). BigWig files generated from depth-normalised, input-controlled datasets were visualised in IGV to assess enrichment at selected genomic loci.

### Protein extraction and western blotting

Bone tissue was pulverised in liquid nitrogen and lysed in RIPA buffer (50 mM Tris–HCl pH 7.5, 150 mM NaCl, 5 mM EDTA, 1% NP-40, 0.5% deoxycholate, 0.1% SDS) with protease/phosphatase inhibitors (Cell Signalling, 5872); clarified at 13,000 rpm (15 min, 4°C). Protein was quantified by BCA (Thermo Fisher). Equal amounts (20 μg osteoblasts; 40 μg bone) were resolved by SDS-PAGE (8–15%) and transferred to nitrocellulose (Amersham). Membranes were blocked in 2.5% BSA/TBST; primary antibodies (Table S3) were applied overnight at 4°C, followed by HRP-conjugated secondary antibodies (1:10,000, 1 h RT). Signal was detected with ECL (BioRad) on an Azure 300 imager and quantified with ImageJ. For phosphorylation analyses, phosphoprotein signals were normalized to the corresponding total protein signal from the same sample. For total protein analyses, target protein signals were normalized to β- actin.

### Staining assays

For Alkaline phosphatase (ALP) staining, primary osteoblasts were differentiated for 7 days. Cells were washed with PBS, fixed with 4% PFA and incubated with BCIP/NBT substrate solution (Sigma-Aldrich, B1911) in the dark at 37°C. Staining was visualised using an inverted light microscope, and four non-overlapping fields were imaged per sample. Mineralization was assessed after 20 days of osteogenic differentiation by Alizarin Red S (ARS) staining. Cells were fixed with 4% PFA, stained with 2% Alizarin Red S solution (Sigma, TMS-008-C) and washed extensively with water for excess dye removal. For quantification, bound dye was extracted with 10% acetic acid, heated (85°C, 10 min), centrifuged (12,000 × g, 10 min), neutralised with 1 M NaOH, and absorbance measured at 405 nm. For image-based quantification, ARS-stained cultures were imaged under identical conditions, and the stained area was quantified using ImageJ and expressed as a percentage of the total area of each microscopic field. Multiple fields were analyzed per sample and averaged to obtain a single value for each sample. Sirius Red staining: cell monolayers were fixed in 3.7% formaldehyde, stained with 0.1% Direct Red 80 in saturated picric acid (pH 3.5, 1 h), and washed with 0.01 N HCl. The stained area was quantified using ImageJ and expressed as a percentage of the total area of each microscopic field. Multiple fields were analyzed per sample and averaged to obtain a single value for each sample.

### Live-cell calcium imaging

Spontaneous Ca² transients were recorded with Fluo-4 AM (4 μM with 0.04% Pluronic F-127, 45 min at 37°C; 20 min de-esterification) on a Zeiss LSM 980 confocal microscope at 37°C, with identical acquisition settings across all groups. Day 4 was selected as an intermediate stage of osteoblast differentiation to assess intracellular Ca² dynamics before the late-stage mineralisation phenotype becomes apparent. Images were acquired at 3 s intervals for 72 s (24 frames). F was defined as the mean intensity of the five quietest consecutive frames per cell; ΔF/F = (F − F )/F . Mean, peak, and SD of ΔF/F were calculated per cell.

### Seahorse metabolic assays

OCR was measured with the Cell Mito Stress Test and PER with the Glycolytic Rate Assay on a Seahorse XF24 analyser (Agilent), with cells (5 × 10 /well) analysed at day 0 (undifferentiated) or day 7 of differentiation, the latest time point at which primary osteoblast cultures could be reliably maintained on Seahorse plates for extracellular flux measurements. These time points enabled comparison of the basal metabolic state with metabolic changes during osteoblast differentiation, before the overt late-stage mineralisation phenotype. Mito Stress: base medium (2 mM glutamine, 1 mM pyruvate, 5.5 mM glucose, pH 7.4); sequential injection of oligomycin (1 μM), FCCP (2 μM), and rotenone/antimycin A (0.5 μM each). Glycolytic Rate: base medium (10 mM glucose); rotenone/antimycin A then 2-DG (50 mM). Both assays used 1 h CO -free equilibration at 37°C. Data acquired with Seahorse Wave v2.6.3.5.

### Mitochondrial superoxide measurement

Osteoblasts were stained with 200 nM MitoTracker Deep Red FM (Invitrogen, 30 min, 37°C) followed by 2.5 μM MitoSOX Red (Invitrogen, 10 min, protected from light) in serum-free medium, washed with PBS, and immediately imaged on a Zeiss LSM 980. MitoTracker-positive structures were segmented; median MitoSOX fluorescence within mitochondrial ROIs was quantified in ImageJ across three independent biological replicates.

### Serum metabolomics

Untargeted metabolomics: Serum samples from six mice per group (n = 6) was extracted with ice-cold methanol (14), centrifuged, dried, and reconstituted with internal standards for UHPLC–Q Exactive Plus MS analysis (Thermo Fisher). Raw data were processed with Compound Discoverer and MetaboAnalyst 5.0(15); FDR < 0.1, |log FC| > 0.5). Targeted GLA quantification: Serum samples from 4–5 mice per group (n = 4–5) were subjected to lipid extraction by modified Folch method (16), resuspended in 200 μL 2:1 chloroform:methanol, and analysed by LC-MS on an Agilent 6545 Q-TOF in negative ionisation mode (30 min, dual AJS-ESI); quantification was by AUC normalised to C15:0 free fatty acid internal standard and serum volume.

### Statistical analysis

Statistical analysis was performed with GraphPad Prism 10. Two-group comparisons used unpaired two-tailed Student’s t-tests; multiple groups used one-way or two-way ANOVA with Tukey’s post hoc test as indicated. Data are presented as mean ± SEM. The unit of analysis is a single mouse and the number of biological replicates are indicated as individual data points. *P* values for relevant comparisons are indicated on the plots.

### Data availability

The transcriptomics data generated in this study have been deposited in the NCBI Gene Expression Omnibus (GEO) under accession number GSE336941. The data supporting the findings of this study are provided in the article and its Supporting Information.

### Supporting information

This article contains supporting information.

## Acknowledgements

We are grateful to Mr. Khem Singh Negi for technical support, Mr. B. N. Roy for histopathology assistance, Dr. Nagarajan and the Small Animal Facility staff for their help with animal experiments, and the Confocal Microscopy and Next-Generation Sequencing facilities at NII for technical support with imaging and sequencing. The authors declare no conflict of interest.

## Conflict of interest

The authors declare no conflicts of interest regarding the content of this article.

## Funding

GAA acknowledges intramural research funds of the National Institute of Immunology and the Department of Biotechnology, Govt. of India (grant numbers BT/PR45456/PFN/20/1596/2022 and BT/PR48040/PFN/20/1617/2022). SSK acknowledges the SwarnaJayanti Fellowship from the Anusandhan National Research Foundation, Government of India (grant number: SB/SJF/2021-22/01).

## Abbreviations

ALP,: alkaline phosphatase;
ARS,: Alizarin Red S;
BMP,: bone morphogenetic protein;
CaMKII,: Ca² /calmodulin-dependent protein kinase II;
DMSO,: dimethyl sulfoxide;
ECM,: extracellular matrix;
ECL,: enhanced chemiluminescence;
FBS,: fetal bovine serum;
FDR,: false discovery rate;
GLA,: γ-linolenic acid;
GO,: Gene Ontology;
H&E,: hematoxylin and eosin;
LC-MS,: liquid chromatography–mass spectrometry;
MBD,: milk-based diet;
OCR,: oxygen consumption rate;
PFA,: paraformaldehyde;
PPAR,: peroxisome proliferator-activated receptor;
qPCR,: quantitative PCR;
RIPA,: radioimmunoprecipitation assay;
ROS,: reactive oxygen species;
RXR,: retinoid X receptor;
SEM,: standard error of the mean;
SOCE,: store-operated calcium entry;
TGF-β,: transforming growth factor-β;
UHPLC,: ultra-high-performance liquid chromatography;
VDR,: vitamin D receptor;
WT: wild type

The authors have declared that no conflict of interest exists.

---

**Figure S1.**
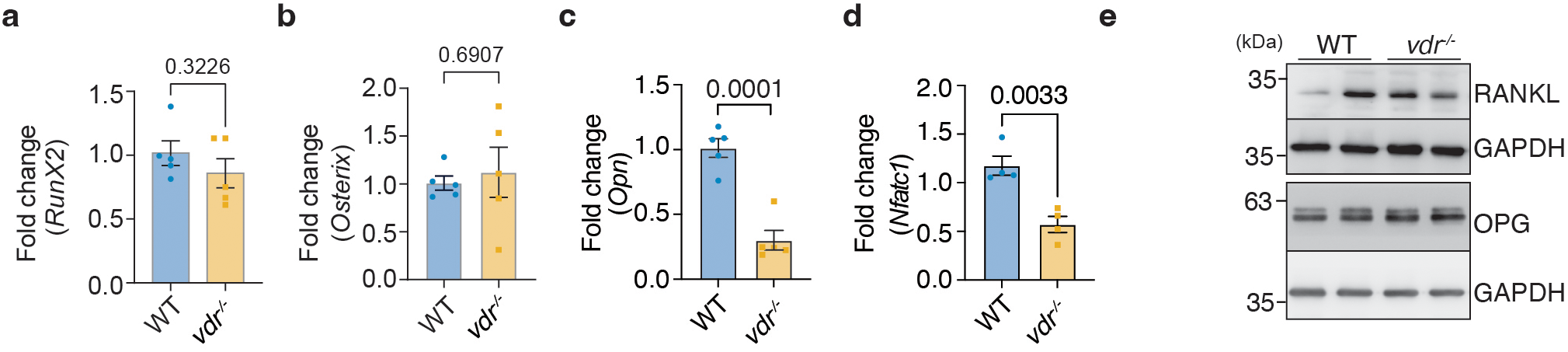
Expression analysis of osteogenic and osteoclast-associated genes in WT and *vdr /* bones. (a-b) Quantitative PCR analysis of *Runx2* and *Sp7* (Osterix) expression in bone tissue from WT and *vdr /* mice. (c-d) Quantitative PCR analysis of Opn and Nfatc1 expression in bone tissue from WT and *vdr /* mice. (e) Western blot analysis of Rankl and Opg expression in bone tissue from WT and *vdr /*mice. Data are presented as mean ± SEM. Statistical analyses were performed using an unpaired two-tailed Student’s t-test. *P < 0.05, **P < 0.01, ***P < 0.001.Individual biological replicates are denoted by dots on the graphs, with the number of biological replicates (*n*) indicated in each graph.

**Figure S2.**
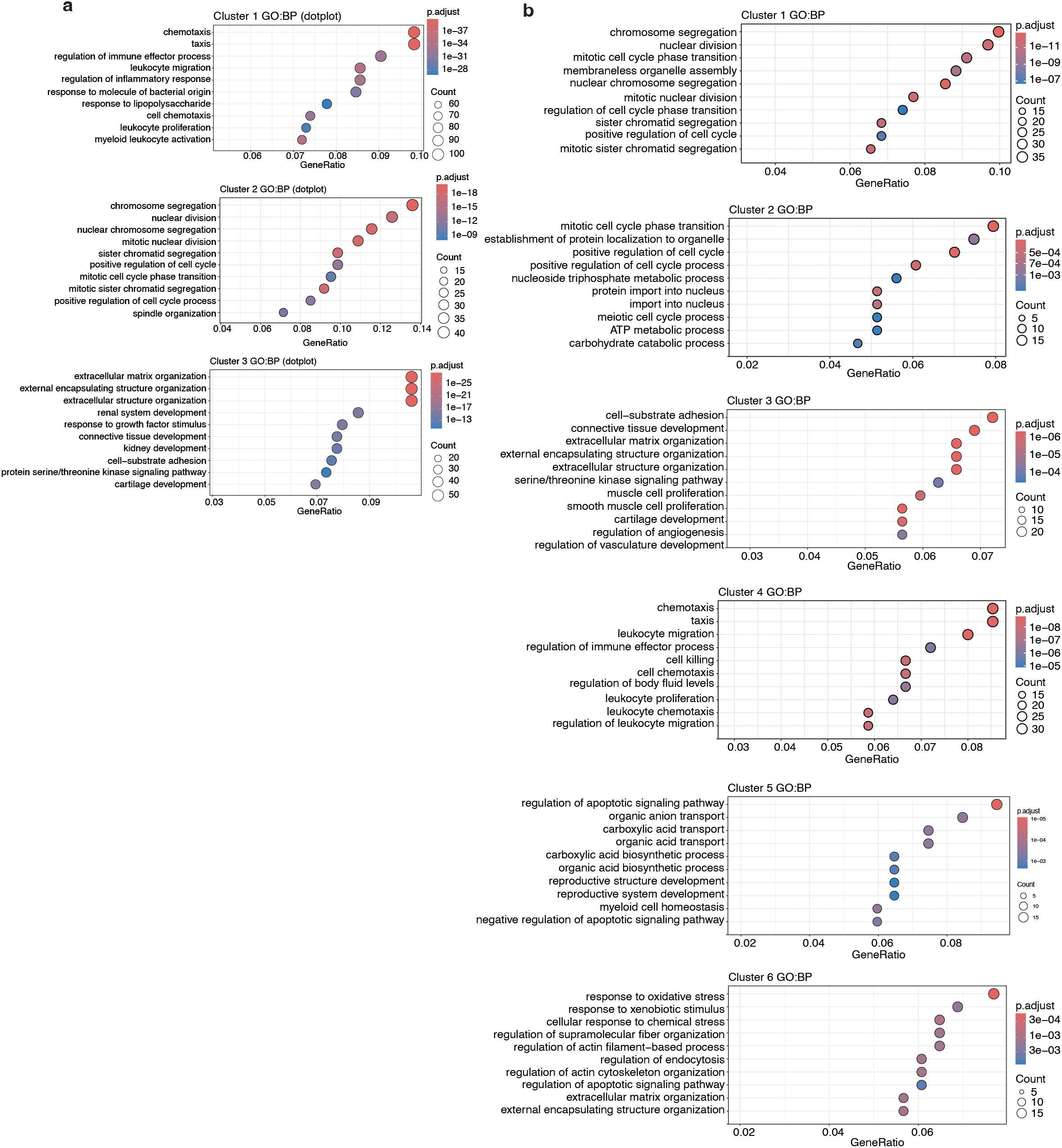
Transcriptomic clustering and pathway enrichment analysis during osteoblast differentiation in WT and *vdr /* osteoblasts. (a) Gene ontology (GO) enrichment analysis for clusters identified in Figure 3a. (b) GO enrichment analysis of clusters identified in the clustering analysis of gene expression data shown in Figure 3b.

**Figure S3.**
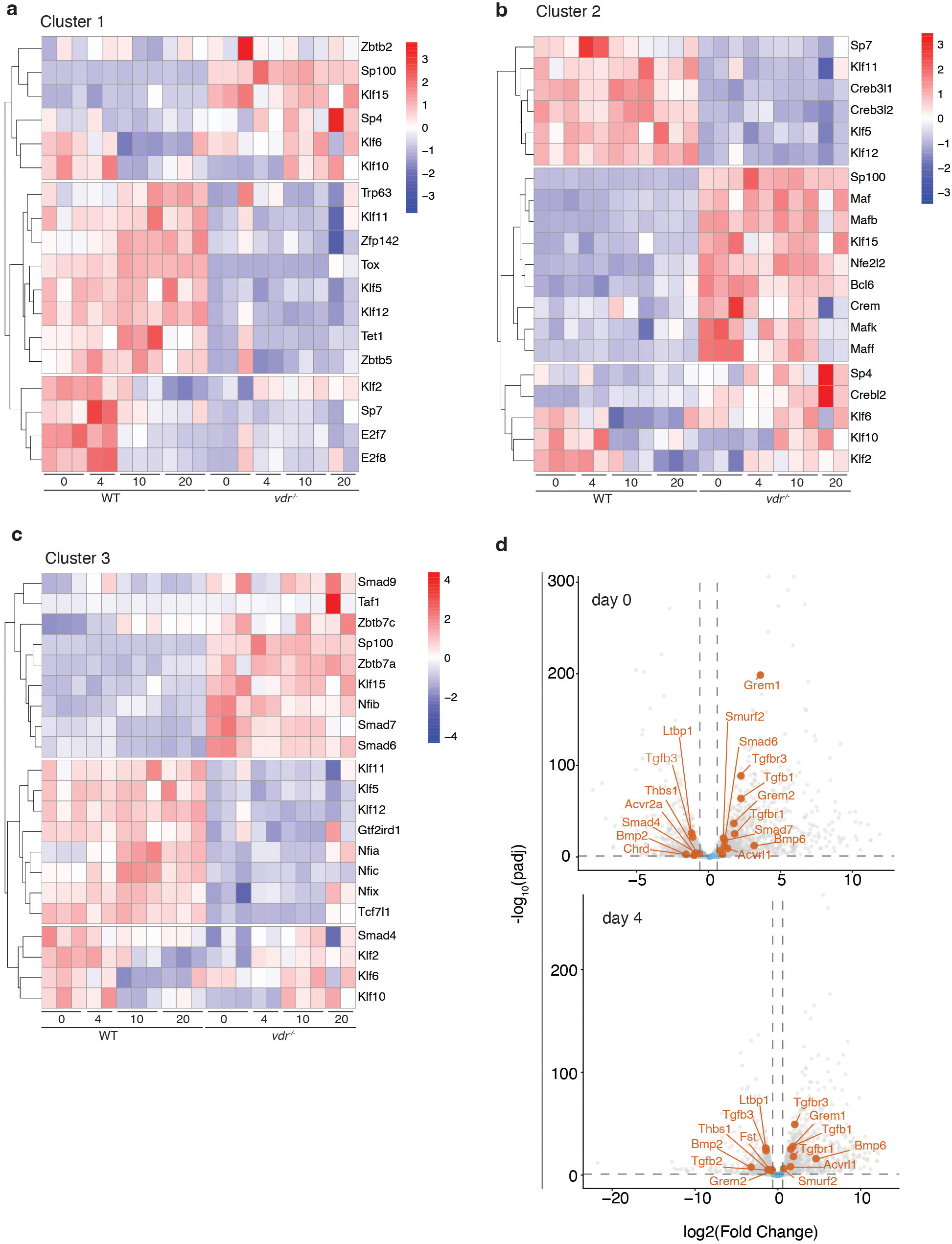
Transcription factor enrichment analysis of differentiation-associated gene clusters in WT and *vdr /* osteoblasts. (a-c) Transcription factor enrichment analysis of genes associated with clusters 1, 2, and 3 identified during osteoblast differentiation. Transcription factors whose binding sites are enriched on the promoters of genes belonging to the clusters identified in Figure 3b were identified using TRANSFAC. (d) Volcano plots of differentially expressed genes at days 0 and 4 of differentiation, highlighting altered expression of BMP/TGF-β pathway components in *vdr /* osteoblasts.

**Figure S4.**
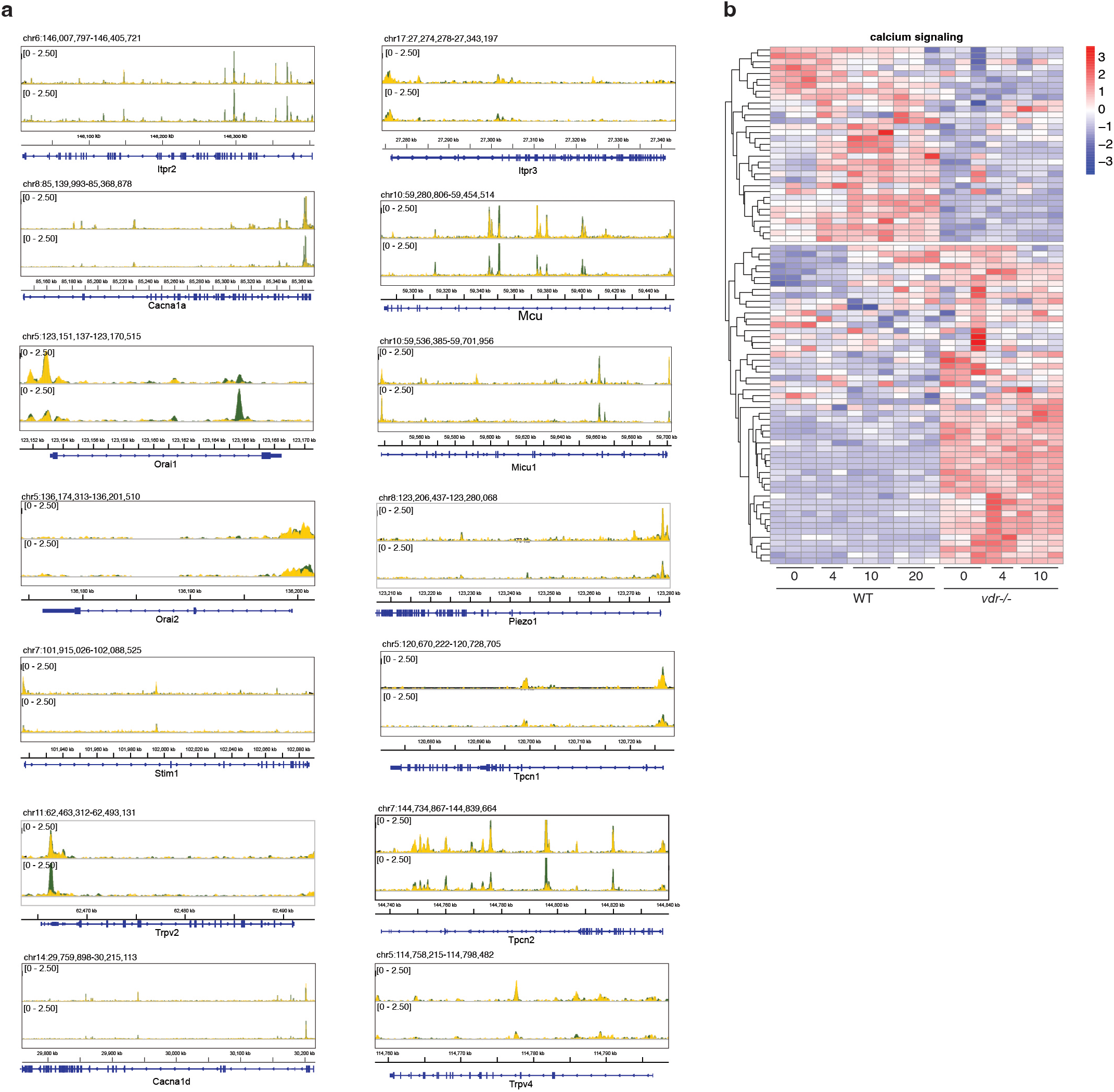
VDR deficiency impairs intracellular calcium signalling and CaMKII activation in osteoblasts. (a) ChIP-seq tracks showing RXR (top) and VDR (bottom) occupancy across the genomic loci of selected Ca² channel and regulatory genes in osteoblasts treated with vehicle (yellow) or 1,25(OH) D (green). Each panel displays chromosomal coordinates (mm10) and the corresponding gene model (blue) beneath paired ChIP-seq signal tracks scaled to [0–2.50]. (b) Heatmap showing expression of genes associated with calcium transport, intracellular calcium homeostasis, and organellar calcium signalling in WT and *vdr /* osteoblasts during differentiation.

**Figure S5.**
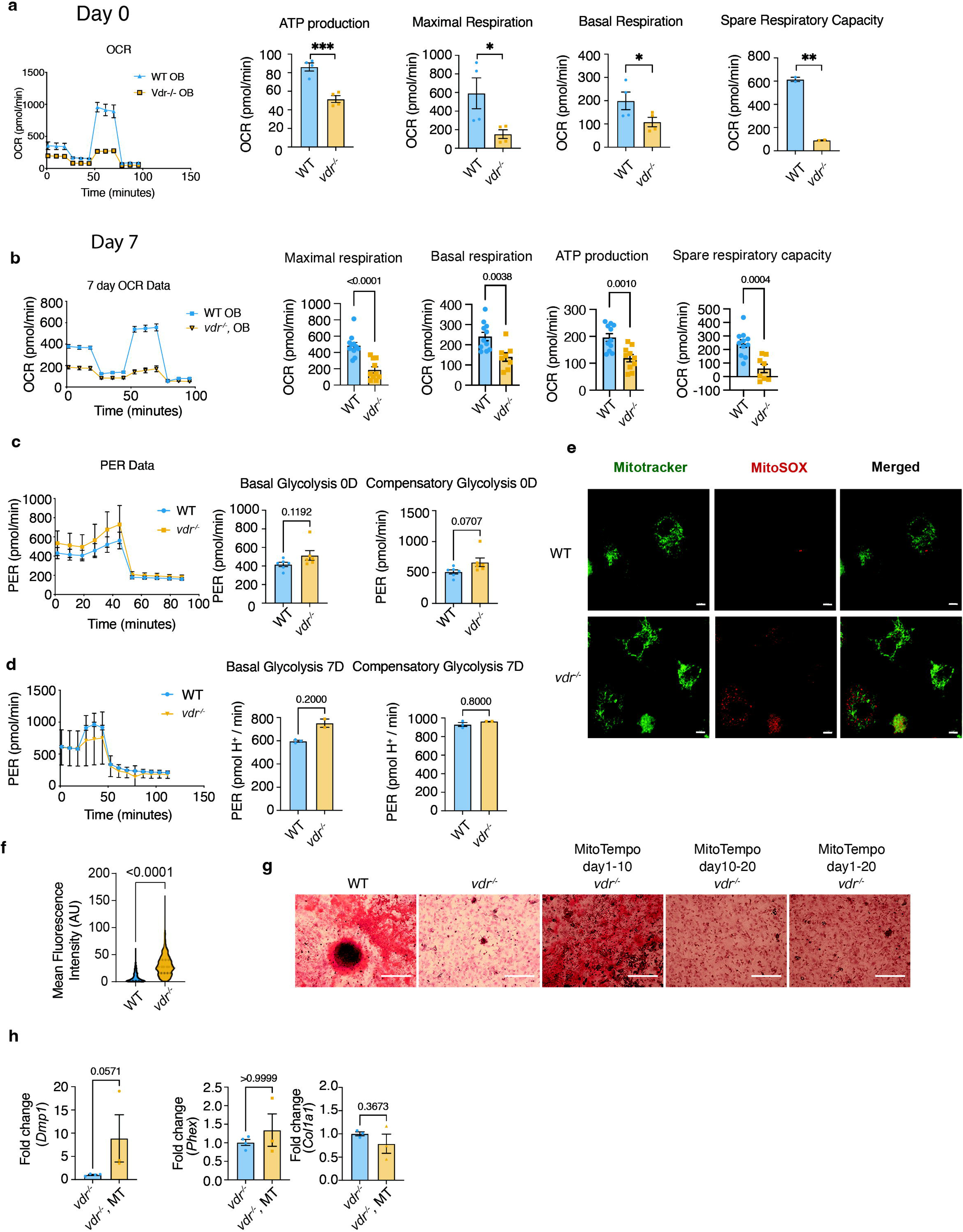
*vdr /* osteoblasts exhibit mitochondrial dysfunction and altered redox homeostasis. (a-b) Seahorse XF analysis of oxygen consumption rate (OCR) parameters in WT and *vdr /*osteoblasts. (c-d) Seahorse XF analysis of proton efflux rate (PER) in WT and *vdr /*osteoblasts. (e) Representative MitoTracker and MitoSOX staining for mitochondrial ROS in WT and *vdr /* osteoblasts on day 7 of differentiation. (f) Quantification of MitoSOX signals in WT and *vdr /* osteoblasts shown in (e). (g) Representative Alizarin Red staining of WT and *vdr /* osteoblast cultures following MitoTEMPO treatment (Scale bar= 300μm). (h) Quantitative PCR analysis of osteogenic marker expression in WT and *vdr /* osteoblasts following MitoTEMPO treatment. For figure f, data are shown as violin plots (median ± quartiles) and analyzed by a Mann-Whitney test. All other figures are presented as mean ± SEM and analyzed via unpaired two-tailed Student’s t-test where individual biological replicates are denoted by dots on the graphs, with the number of biological replicates (*n*) indicated in each graph. *P < 0.05, **P < 0.01, ***P < 0.001.

**Figure S6.**
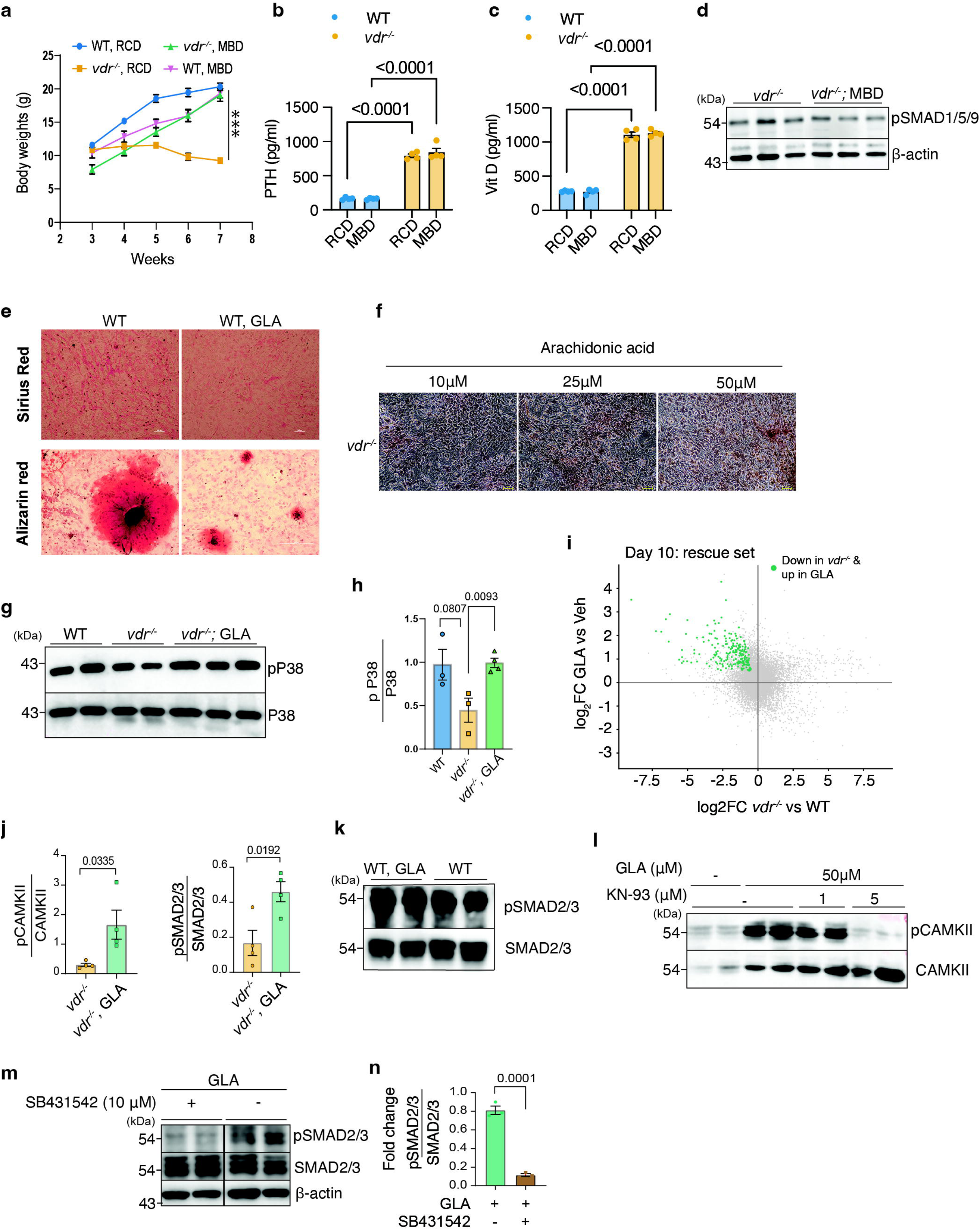
Milk-based diet and **γ**-linolenic acid restore osteoblast maturation-associated defects in *vdr /* mice and osteoblasts. **(a)** Body weight of WT and *vdr* / mice maintained on regular chow diet (RCD) or milk-based diet (MBD) from the indicated ages. **(b)** Serum parathyroid hormone (PTH) concentrations in WT and *vdr* / mice maintained on RCD or MBD. **(c)** Serum 25-hydroxyvitamin D [25(OH)D] concentrations in WT and *vdr* / mice maintained on RCD or MBD. **(d)** Representative immunoblot showing phospho-SMAD1/5/9 and β-actin in bone tissue from *vdr* / mice maintained on RCD or MBD. **(e)** Representative Sirius Red and Alizarin Red staining of WT osteoblast cultures treated with vehicle or 50 μM GLA. **(f)** Representative Alizarin Red staining of *vdr* / osteoblast cultures treated with the indicated concentrations of arachidonic acid (scale bar= 48μm). **(g)** Representative immunoblot showing phospho-p38 and total p38 in WT, untreated *vdr* / , and GLA-treated *vdr* / osteoblasts at day 20 of differentiation. **(h)** Quantification of phospho-p38 relative to total p38. **(i)** Scatter plot showing transcriptomic changes in *vdr* / osteoblasts relative to WT (x-axis) and following GLA treatment of *vdr* / osteoblasts (y-axis). The GLA rescue set comprises genes significantly downregulated in *vdr* / osteoblasts relative to WT and significantly upregulated following GLA treatment; these genes are highlighted in green. **(j)** Quantification of phospho-CaMKII relative to total CaMKII and phospho-SMAD2/3 relative to total SMAD2/3 in untreated and GLA-treated *vdr* / osteoblasts. **(k)** Representative immunoblot showing phospho-SMAD2/3 and total SMAD2/3 in WT osteoblasts treated with 50 μM GLA or vehicle. **(l)** Representative immunoblot showing phospho-CaMKII and total CaMKII in *vdr* / osteoblasts treated with GLA (50 μM), KN-93 (1 or 5 μM), or the indicated combinations. **(m)** Representative immunoblot showing phospho-SMAD2/3, total SMAD2/3, and β-actin in GLA-treated *vdr* / osteoblasts ± SB431542 (10 μM). The blot was spliced as indicated by the vertical line. **(n)** Quantification of phospho-SMAD2/3 relative to total SMAD2/3 following SB431542 treatment. Data are presented as mean ± SEM. For multi-group comparisons (panels a–c), data were analyzed by two-way ANOVA followed by Tukey’s post hoc test. For panels h,j , and n, comparisons were analyzed using an unpaired two-tailed Student’s *t*-test. *P* < 0.05, \**P* < 0.01, \*\**P* < 0.001. Individual biological replicates are denoted by dots on the graphs, with the number of biological replicates (*n*) indicated in each graph.

## Notes

### Competing Interest Statement

The authors have declared no competing interest.

### Summary of Updates

The manuscript size was reduced. WT-GLA experiments were added.

