## Supplemental Tables for "γ-Linolenic Acid Induces a Vitamin D Receptor-Independent Mineralization Program by Activating CaMKII–SMAD2/3 Pathway in Calvarial Osteoblasts"

**Table S1**

| **Sl. No** | **Probe ID** | **Accession** | **Target Sequence** | **HUGO Gene Symbol** | **NSID** |
| --- | --- | --- | --- | --- | --- |
| 1 | ACC | NM_133904.2 | CCTGCTGATTCTCTGCTGAACATTGTGGACGTTGAATTGATTTACGGCGGCATCAAGTATGCTCTCAAGGTAGCCCGACAGTCCCTGACTATGTTTGTCC | Acacb | NM_133904.2:2382 |
| 2 | MCAD | NM_007382.4 | AAGGAAATGAGATCAAAGACGAGTTTGGATCTGTGCAGCGGATTCCCATGGCGGAGGAACCTGTCTTCAGCTCTATGGTGACCCTTTCTAGATAGGTTTG | Acadm | NM_007382.4:1365 |
| 3 | SCAD | NM_007383.2 | GTGTTCGGGACTGGCGACGGTTACACACTGTTTACCAGTCTGTGGAGCTGCCTGAGACACACCAGATGTTGCGTCAGACATGCCGTGACTTTGCCGAGAA | Acads | NM_007383.2:95 |
| 4 | VLCAD | NM_017366.2 | ACATTTTCACTGTCTTTGCCAAGACGCCAATTAAAGATGCAGCCACGGGGGCCGTGAAAGAGAAGATCACAGCTTTTGTAGTGGAAAGGAGCTTCGGAGG | Acadvl | NM_017366.2:855 |
| 5 | ACS | NM_019811.3 | AGATTAGAGAAAAGATTGGCCCCATTGCCACACCAGACTACATCCAGAATGCACCTGGCTTGCCTAAAACACGCTCAGGGAAAATCATGAGGCGAGTTCT | Acss2 | NM_019811.3:1964 |
| 6 | ALP | NM_007431.2 | CTCCAGCGAGGGACGAATCTCAGGGTACACCATGATCTCACCATTTTTAGTACTGGCCATCGGCACCTGCCTTACCAACTCTTTTGTGCCAGAGAAAGAG | Alpl | NM_007431.2:205 |
| 7 | ANK | NM_020332.4 | CCTCTTGGTCAACGTTTCTACTCCCTTGGACAATCTCCACTTTGGAACCAAAGGACTTGGGCCAGACTTTTCCTGTTCATGTTTGCCTCCTCCTAAGAAT | Ank | NM_020332.4:2075 |
| 8 | PMCA1b | NM_026482.1 | AAACAAGTCATGGGCCAGTGGTCAAGATACTTCTCTGGGAATTGTTGCTGCTGATACTGCTCTTCAGAGTCCTACAATGAGTGTGTGGTTTAAGGAAGAT | Atp2b1 | NM_026482.1:230 |
| 9 | BDNF | NM_007540.4 | AGTCCCGTCTGTACTTTACCCTTTGGGGTTAGAAGTCAAGTTGGAAGCCTGAATGAATGGACCCAATGAGAACTAGTGTTAAGCCCATTTCCCTAGTCAG | Bdnf | NM_007540.4:3260 |
| 10 | Osteocalcin | NM_007541.2 | GTGAGCTTAACCCTGCTTGTGACGAGCTATCAGACCAGTATGGCTTGAAGACCGCCTACAAACGCATCTATGGTATCACTATTTAGGACCTGTGCTGCCC | Bglap | NM_007541.2:251 |
| 11 | BMP2 | NM_007553.2 | GAGGATTAGCAGGTCTTTGCACCAAGATGAACACAGCTGGTCACAGATAAGGCCATTGCTAGTGACTTTTGGACATGATGGAAAAGGACATCCGCTCCAC | Bmp2 | NM_007553.2:885 |
| 12 | BMP4 | NM_007554.2 | AGACCCTAGTCAACTCTGTTAATTCTAGTATCCCTAAGGCCTGTTGTGTCCCCACTGAACTGAGTGCCATTTCCATGTTGTACCTGGATGAGTATGACAA | Bmp4 | NM_007554.2:1500 |
| 13 | BMP7 | NM_007557.2 | GGTTTGATCTTTCCAAGATCCCCGAGGGCGAAGCGGTGACCGCAGCCGAATTCAGGATCTATAAGGACTACATCCGGGAGCGATTTGACAACGAGACCTT | Bmp7 | NM_007557.2:690 |
| 14 | CD36 | NM_007643.3 | GGGACCATTGGTGATGAAAAAGCAGAAATGTTCAAAACACAAGTGACTGGGAAAATCAAGCTCCTTGGCATGGTAGAGATGGCCTTACTTGGGATTGGAG | Cd36 | NM_007643.3:1520 |
| 15 | CNTF | NM_170786.2 | ATGGGGGTGCACAATCCCATTAGAGAATGCCCAGTAGATTTAGTCTGTGGGAGTCACATTTCTTATTTGGACTAGTGAAGACAGAAGCAAACCAGCTCAC | Cntf | NM_170786.2:110 |
| 16 | CPT1b | NM_009948.2 | ACAAGATGTCTCTGGACGCCATCGAACGTGCTGCTTTCTTTGTGACCCTGGATGAAGATTCTCATTGCTACAACCCTGACGATGAGACCAGTCTTAGCCT | Cpt1b | NM_009948.2:1297 |
| 17 | CPT2 | NM_009949.2 | GAGTTTCTCCACTGTGTCCAGAAGTGCTTGGAAGACATGTTCGATGCCCTCGAAGGCAAAGCCATCAAAACTTAGCTTCTTGGTCGATGAAAAGCCTCCA | Cpt2 | NM_009949.2:2025 |
| 18 | Decorin | NM_001190451.1 | CCAAACACCTTCAGATGTGTCTATGTGCGTTCTGCCATTCAACTTGGAAACTACAAGTAACCCTCAGACGGCCTAATTCTTATAATCTGGAAAAACACCC | Dcn | NM_001190451.1:1319 |
| 19 | DMP1 | NM_016779.2 | AAGACTGTCATTCTCCTTGTGTTCCTTTGGGGGCTGTCCTGTGCTCTCCCAGTTGCCAGATACCACAATACTGAATCTGAAAGCTCTGAAGAGAGGACGG | Dmp1 | NM_016779.2:120 |
| 20 | DRP1 | NM_001025947.1 | GGTTAATCATGTGAAAGATACTCTTCAGAGTGAACTGGTAGGGCAGCTGTATAAGTCATCCTTATTAGATGACCTTCTGACTGAATCCGAGGACATGGCC | Dnm1l | NM_001025947.1:2075 |
| 21 | ENPP1 | NM_008813.3 | GTGGAAGTGGATTTCATGGCTCTGACAACTTGTTTTCAAACATGCAAGCTCTTTTTATTGGCTATGGACCTGCCTTCAAGCATGGTGCTGAAGTTGACTC | Enpp1 | NM_008813.3:1645 |
| 22 | FABP3 | NM_010174.1 | GAACGGGGATACTATCACCATAAAGACACAAAGTACCTTCAAGAACACAGAGATCAACTTTCAGCTGGGAATAGAGTTCGACGAGGTGACAGCAGATGAC | Fabp3 | NM_010174.1:134 |
| 23 | FABP4 | NM_024406.2 | TCGAAGGTTTACAAAATGTGTGATGCCTTTGTGGGAACCTGGAAGCTTGTCTCCAGTGAAAACTTCGATGATTACATGAAAGAAGTGGGAGTGGGCTTTG | Fabp4 | NM_024406.2:50 |
| 24 | FGF2 | NM_008006.2 | CTCTACTGCAAGAACGGCGGCTTCTTCCTGCGCATCCATCCCGACGGCCGCGTGGATGGCGTCCGCGAGAAGAGCGACCCACACGTCAAACTACAACTCC | Fgf2 | NM_008006.2:287 |
| 25 | FGF 23 | NM_022657.4 | TCTCCACGGCAACATTTTTGGATCGCTTCACTTCAGCCCAGAGAATTGCAAGTTCCGCCAGTGGACGCTGGAGAATGGCTATGACGTCTACTTGTCGCAG | Fgf23 | NM_022657.4:418 |
| 26 | Irisin | NM_027402.3 | TACCTTACATGAAATCCATTCAACCGGCTCTGGACACTTCTCTTATCTGTTCCCTTCCTGGTTTCCAAAGAAGCCCTCTGTGAACATCATCAAAGCATGG | Fndc5 | NM_027402.3:1142 |
| 27 | HMBS | NM_013551.2 | CGTTCACTCCCTGAAGGATGTGCCTACCATACTACCTCCTGGCTTTACTATTGGAGCCATCTGCAAACGGGAAAACCCTTGTGATGCTGTTGTCTTTCAC | Hmbs | NM_013551.2:464 |
| 28 | HPRT1 | NM_013556.2 | TGCTGAGGCGGCGAGGGAGAGCGTTGGGCTTACCTCACTGCTTTCCGGAGCGGTAGCACCTCCTCCGCCGGCTTCCTCCTCAGACCGCTTTTTGCCGCGA | Hprt | NM_013556.2:30 |
| 29 | IGF1 | NM_001111274.1 | GGGCTTTTACTTCAACAAGCCCACAGGCTATGGCTCCAGCATTCGGAGGGCACCTCAGACAGGCATTGTGGATGAGTGTTGCTTCCGGAGCTGTGATCTG | Igf1 | NM_001111274.1:418 |
| 30 | IL15 | NM_008357.2 | CTTGCAAACAGCACTCTGTCTTCTAACAAGAATGTAGCAGAATCTGGCTGCAAGGAATGTGAGGAGCTGGAGGAGAAAACCTTCACAGAGTTTTTGCAAA | Il15 | NM_008357.2:854 |
| 31 | IL6 | NM_031168.1 | CTCTCTGCAAGAGACTTCCATCCAGTTGCCTTCTTGGGACTGATGCTGGTGACAACCACGGCCTTCCCTACTTCACAAGTCCGGAGAGGAGACTTCACAG | Il6 | NM_031168.1:40 |
| 32 | IL7 | NM_008371.2 | AAACATTCATTGGTGAACCACTGGGGGAGTGGAACTGTCCTGTTTTAGACTGGAGATACTGGAGGGCTCACGGTGATGGATAATGCTCTTGAAAACAAGA | Il7 | NM_008371.2:1055 |
| 33 | LIF | NM_008501.2 | ACATCTTTCACCTGGAAGCATTGACTTCCACCGAGCATAGTAGGTAGTGTGTCTGGACCAGAGAAAAAGGGATGGGGCATTTTGCAGTTTATCCAGAGAG | Lif | NM_008501.2:3435 |
| 34 | Osteoglycin | NM_008760.4 | GAAATGAAAACAAAGTCTACACCATTGTGCTGTGTGCAGCTTCACCTGTATATGTTCACCCAAACAAACTAAATATGCTCATTCCAGCCCAAGAGCACCA | Ogn | NM_008760.4:1795 |
| 35 | Polr2b | NM_153798.2 | CGGTTCAAGAGGGGCAAAGCCTGGTGTTACTAAGGAGAAAAGAATTAAATATGCAAAAGAAGTCTTACAGAAAGAAATGCTCCCTCACGTTGGTGTCAGT | Polr2b | NM_153798.2:1090 |
| 36 | PTGES | NM_022415.2 | GAAGGGAACTTTGGCTTCTTCAGCATCTGTGAGGTTTGAAGATGCCAAATGTCTACTCCTGTGGAAAATCTCACTTGTGTGTGTGCAGTGTCCCTCCAGA | Ptges | NM_022415.2:1185 |
| 37 | CACT | NM_020520.4 | TGCGCAAAGAAGCTGTATCAGGAGTTCGGGATCCGCGGCTTCTACAAAGGGACTGTGCTCACACTCATGCGAGATGTTCCTGCCAGTGGGATGTATTTCA | Slc25a20 | NM_020520.4:592 |
| 38 | FATP1/ SLC27A1 | NM_011977.3 | CTCTGCGGCGTTTCGATGGTTATGTTAGTGACAGTGCCACCAACAAGAAGATTGCCCACAGCGTTTTCCGAAAGGGCGATAGCGCCTACCTCTCAGGTGA | Slc27a1 | NM_011977.3:1580 |
| 39 | FATP4 | NM_011989.4 | TAGCCCCTTTCGCTTAGCCCTTGGGATAAGCCCTTAGCCAGTCCTCTTTTCCAAAGCTCTGCCTTCCTATAACTCCTTAGGAGAGACTCCCACCAGATCT | Slc27a4 | NM_011989.4:2980 |
| 40 | slc8a1 | NM_011406.3 | TAAGTCTCCCACCCAATGTTTCAATGGGATTTCGTCTGGTAGCTCTGGTGGCTCTCTTGTTTTCCCATGTTGACCATATAACTGCAGATACAGAGGCAGA | Slc8a1 | NM_011406.3:120 |
| 41 | SMPD3 | NM_021491.3 | TAAGTTGAAAGAGCAGCTACACGGCTACTTCGAGTACATCCTGTATGATGTTGGGGTCTACGGTTGTCATGGTTGCTGCAATTTCAAATGTCTCAACAGC | Smpd3 | NM_021491.3:1578 |
| 42 | SOST | NM_024449.4 | ACGAAAGACCTGGGACTGGTTATGGACGTACAGTAAGATCTACTCCTTCCACCCAAATGTAAAGCCTGCGTGGGCTAGATAGGGTTTCTGACCCTGACCT | Sost | NM_024449.4:975 |
| 43 | Osteopontin | NM_009263.3 | TGAATCTGACGAATCTCACCATTCGGATGAGTCTGATGAGACCGTCACTGCTAGTACACAAGCAGACACTTTCACTCCAATCGTCCCTACAGTCGATGTC | Spp1 | NM_009263.3:420 |
| 44 | TBP | NM_013684.3 | GTGGCGGGTATCTGCTGGCGGTTTGGCTAGGTTTCTGCGGTCGCGTCATTTTCTCCGCAGTGCCCAGCATCACTATTTCATGGTGTGTGAAGATAACCCA | Tbp | NM_013684.3:70 |
| 45 | TGF beta1 | NM_011577.1 | GGAGTTGTACGGCAGTGGCTGAACCAAGGAGACGGAATACAGGGCTTTCGATTCAGCGCTCACTGCTCTTGTGACAGCAAAGATAACAAACTCCACGTGG | Tgfb1 | NM_011577.1:1470 |
| 46 | TNF alpha | NM_013693.2 | TGGATCTCAAAGACAACCAACTAGTGGTGCCAGCCGATGGGTTGTACCTTGTCTACTCCCAGGTTCTCTTCAAGGGACAAGGCTGCCCCGACTACGTGCT | Tnf | NM_013693.2:514 |
| 47 | WNT-3a | NM_009522.2 | AGATCCTACCTGTGAGGGTCTCATACCTAAGGACCCGGTTTCTGCCTTCAGCCTGGGCTCCTATTTGGGATCTGGGTTCCTTTTTAGGGGAGAAGCTCCT | Wnt3a | NM_009522.2:1279 |
| 48 | WNT 4 | NM_009523.1 | TGCGGTCCCTGCGACTCCTCGTCTTCGCCGTGTTCTCGGCCGCCGCGAGCAATTGGCTGTACCTGGCCAAGCTGTCATCGGTGGGCAGCATCTCCGAAGA | Wnt4 | NM_009523.1:64 |

Table S1: List of gene names and target sequences of NanoString probes used in figure 1.

**Table S2**

| **Primer** | **Primer length** | **Sequence (5' to 3' direction)** |
| --- | --- | --- |
| Col1a1 qF | 19 | ACTGTCCCAACCCCCAAAG |
| Col1a1 qR | 20 | ACGTATTCTTCCGGGCAGAA |
| Dkk1 qF | 20 | GCCTCCGATCATCAGACGGT |
| Dkk1 qR | 20 | GCAGGTGTGGAGCCTAGAAG |
| Sp7 qF | 23 | CTTCCCAATCCTATTTGCCGTTT |
| Sp7 qR | 24 | CGGCCAGGTTACTAACACCAATCT |
| Runx2 qF | 20 | CCCAGCCACCTTTACCTACA |
| Runx2 qR | 20 | TATGGAGTGCTGCTGGTCTG |
| Sost qF | 22 | TTCCCAGCCCAGTAGAGACCGC |
| Sost qR | 22 | AGAAAGACCCCCATCCCACCCG |
| Dmp1 qF | 20 | TTCGCTGAGGTTTTGACCTT |
| Dmp1 qR | 20 | CCCAAAGGAACACAAGGAGA |
| Mepe qF | 21 | GTCTGTTGGACTGCTCCTCTT |
| Mepe qR | 20 | CACCGTGGGATCAGGATACA |
| Phex qF | 19 | GAAAGGGGACCAACCGAGG |
| Phex qR | 23 | AACTTAGGAGACCTTGACTCACT |
| Ocn qF | 21 | CTGACCTCACAGATCCCAAGC |
| Ocn qR | 21 | TGGTCTGATAGCTCGTCACAA |

Table S2: List of primers used for qRT-PCR analysis

**Table S3:**

| **Western Antibodies** | | | |
| --- | --- | --- | --- |
| **REAGENT or RESOURCE** | **SOURCE** | **IDENTIFIER** | **DILUTION** |
| p38 Antibody | CST | 8690 | 1/1000 |
| phospho-p38 Antibody | CST | 4511 | Jan-00 |
| ERK1/2 Antibody | CST | 9102 | 1/1000 |
| phospho-ERK1/2 Antibody | CST | 9101 | 1/1000 |
| Beta-catenin Antibody | Santa-Cruz | sc-7199 | 1/1000 |
| Osteoproterigin Antibody | Santa-Cruz | sc-390518 | 1/1000 |
| RANKL Antibody | Cloud and Clone | PAA855Mu01 | 1/1000 |
| CAMKII Antibody | CST | 4436 | 1/1000 |
| Phospho-CAMKII Antibody | CST | 12716 | 1/1000 |
| SMAD2/3 Antibody | CST | 5678 | 1/1000 |
| Phospho SMAD2/3 Antibody | CST | 8828 | 1/1000 |
| Beta-actin Antibody | CST | 4967 | 1/1000 |
| Peroxidase AffiniPure™ Goat Anti-Rabbit IgG (H+L) | Jackson ImmunoResearch | AB_2313567 | 1/10000 |

Table S3: List of antibodies used in this study
